# Pickering stabilization of a dynamic intracellular emulsion

**DOI:** 10.1101/2021.06.22.449249

**Authors:** Andrew W. Folkmann, Andrea Putnam, Chiu Fan Lee, Geraldine Seydoux

**Author notes:** These authors contributed equally to this work (Andrew W. Folkmann, Andrea Putnam).

## Abstract

Biomolecular condensates are cellular compartments that form by phase separation in the absence of limiting membranes. Studying the P granules of *C. elegans*, we find that condensate dynamics are regulated by protein clusters that adsorb to the condensate interface. Using *in vitro* reconstitution, live observations and theory, we demonstrate that localized assembly of P granules is controlled by MEG-3, an intrinsically disordered protein that forms low dynamic assemblies on P granules. Following classic Pickering emulsion theory, MEG-3 clusters lower surface tension and slow down coarsening. During zygote polarization, MEG-3 recruits DYRK/MBK-2 kinase to accelerate localized growth of the P granule emulsion. By tuning condensate-cytoplasm exchange, interfacial clusters regulate the structural integrity of biomolecular condensates, reminiscent of the role of lipid bilayers in membrane-bound organelles.

**One Sentence Summary:** Biomolecular condensates are stabilized by interfacial nanoscale protein clusters.

## Main Text

Liquid-liquid phase separation (LLPS) has emerged as a new principle for cellular organization (*1*). Phase separation of proteins, frequently with RNA, creates dense condensates visible by microscopy as micron-scale assemblies containing dozens or more protein and RNA species. Condensates have been reconstituted *in vitro* using purified proteins that interact using multivalent, low-affinity binding sites to generate large, interconnected, and dynamic networks. Some proteins assemble into liquid-like condensates that exhibit fast dynamics where molecules exchange continuously with the dilute phase (*1*). Other proteins form more viscous condensates that become kinetically-arrested over time, in some cases becoming glass-like (*2*, *3*). Fast dynamic condensates form emulsions that coarsen down to a single large condensate (*4*–*7*). In contrast, kinetically-arrested condensates form emulsions that resist coarsening over long time scales (*8*). Like condensates *in vitro*, condensates in cells exhibit a range of dynamic behaviors, but these do not always fit theoretical predictions (*9*, *10*). For example, some condensates resist coarsening for hours, yet maintain fast exchange dynamics and dissolve within minutes in response to changes in the cellular environment (*11*). Several hypotheses have been put forward to explain the lack of coarsening in dynamic cellular emulsions, including physical barriers that keep condensates away from each other (*5*, *12*, *13*), active mechanisms that continuously regenerate small condensates (*14*), and chemical reactions and protein gradients that suppress Ostwald ripening (*14*–*17*) (see Supplemental Text). In this study, we investigate the mechanisms that control the dynamics of P granules, condensates in the *C. elegans* germline. We find that P granule dynamics are controlled by nanoscale protein clusters that adsorb to the condensate interface, a phenomenon first described by Ramsden (1904) and Pickering (1907) for inorganic emulsions (*18*, *19*).

P granules were the first cellular condensates proposed to form by LLPS (*20*). At the core of P granules is a liquid-like phase assembled by PGL proteins (*20*–*22*). During most of the *C. elegans* life cycle, PGL condensates form stable assemblies on the cytoplasmic phase of nuclei (*23*). During the oocyte-to-zygote transition, PGL condensates redistribute to the cytoplasm and undergo two rapid cycles of dissolution and condensation (Fig. 1A). The first cycle occurs during oocyte maturation: most PGL condensates in the maturing oocyte dissolve and reassemble minutes later in the newly fertilized egg (zygote). The second cycle occurs when the zygote becomes polarized along its anterior-posterior axis: PGL condensates in the anterior cytoplasm dissolve while PGL condensates in the posterior grow (Fig. 1A). Factors required for P granule dissolution during oocyte maturation have not yet been identified. Factors required for dissolution during polarization include MEX-5, an RNA-binding protein, MEG-3 and MEG-4, two paralogous intrinsically disordered proteins, and MBK-2, a DYRK family kinase that interacts physically and genetically with MEG-3 (*21*, *24*, *25*). During polarization, MEX-5 becomes enriched in the anterior cytoplasm where it promotes P granule dissolution, possibly by competing with P granule proteins for RNA (*21*, *26*). In zygotes lacking *mbk-2* or *meg-3 meg-4* activity, P granules do not dissolve in the anterior cytoplasm, despite a normal MEX-5 gradient (*27*). In this study, we investigate how the MEGs and MBK-2 collaborate with MEX-5 to regulate P granule dynamics.

**Fig. 1.**
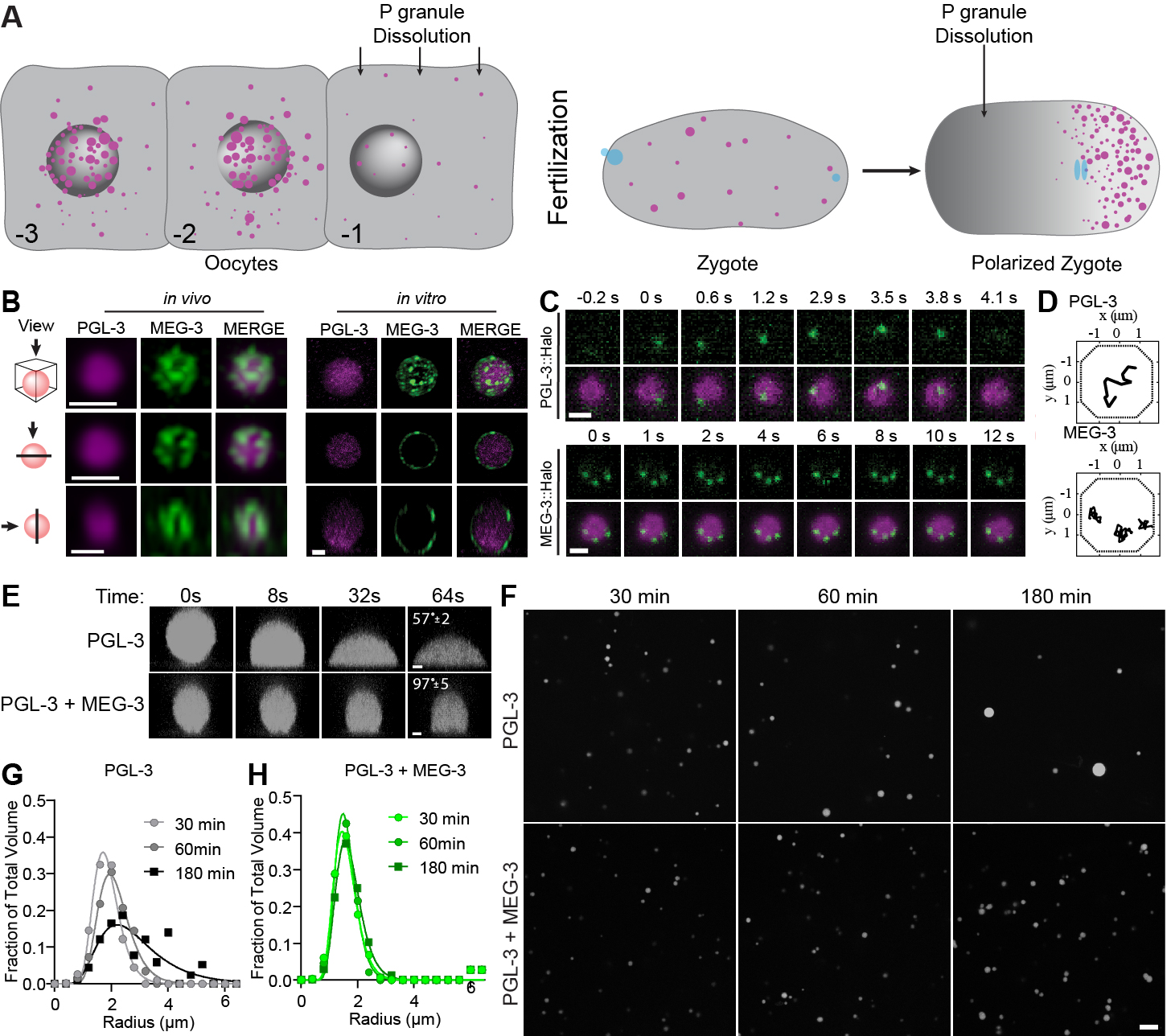
MEG-3 surface-adsorbed clusters reduce surface tension and inhibit coarsening of PGL-3 condensates. A) Schematics depicting the two cycles of P granule dissolution during the oocyte-to-zygote transition. P granules are in magenta. During polarization, the cytoplasmic polarity regulator MEX-5 (gray) becomes enriched in the anterior opposite the P granules. B) Photomicrographs of a P granule *in vivo* labeled with PGL-3::mCherry and MEG-3::meGFP and of a P granule reconstituted *in vitro* PGL-3^488^ and MEG-3^647^. Scale bars are 500 nm and 3 μm respectively. Top panels are a maximum projection of a z-stack through the granule. Middle panels are a single X-Y plane through the middle of the same granule. Lower panels are a single Z-X plane through the middle of the same granule. See fig. S1 for additional examples of P granules captured *in vivo.* C) *In vivo* time lapse series showing single molecules of PGL-3::Halo and MEG-3::Halo (green – JF_646_ dye sparse labeling) in P granules (magenta – JF_549_ dye heavy labeling). Scale bar is 1 μM. D) Representative single molecule trajectories of PGL-3::Halo (top panel) and MEG-3:: Halo (bottom panel) derived from time lapse series as shown in C. Scale bar is 1 μm. See fig. S2 for diffusion rates and dwell times. E) *In vitro* time lapse series showing PGL-3^488^ droplets wetting the glass surface of a glass slide with 3 μM PGL-3, 80 ng/μL *nos-2* RNA, and the presence and absence of 0.5 μM MEG-3 (not shown). Average contact angles at 64s are indicated (See fig. S3A). Scale bar is 1 μm. F) Photomicrographs of a PGL-3^488^ emulsion at the indicated time points after assembly. 3 μM PGL-3 and 80 ng/μL *nos-2* RNA were incubated in condensation buffer in the presence or absence of 0.5 μM MEG-3 (not shown). Scale bar is 5 μm. G and H) Histograms plotting the size distribution of PGL condensates assembled as in (F). Each data point indicates the fraction of total PGL-3 condensate volume represented by condensates binned by radius from 80 images (as in F) collected in 4 independent experiments. Lines fit the data to a log normal distribution.

MEG-3 forms assemblies that accumulate on the surface of PGL-3 condensates in newly fertilized zygotes (*27*, *28*) and in MEG-3/PGL-3 co-condensates reconstituted *in vitro* (*28*). Super resolution 3D confocal microscopy confirmed that MEG-3 forms diffraction-limited clusters (<160 nm) on the PGL-3 interface *in vivo* and *in vitro* (Fig. 1B and fig. S1). MEG-3 clusters are resistant to dilution, high temperature, and salt treatment, in contrast to PGL-3 condensates which readily dissolve in dilute conditions and at elevated temperatures (*28*). Using a single-molecule method adapted from Wu et al., 2019 (*29*), we measured the dynamics of MEG-3 and PGL-3 molecules in P granules *in vivo* (Fig. 1C and D, and movies S1 and S2). Most PGL-3 molecules exhibited short-lived trajectories in P granules with an average apparent diffusion coefficient of *D_c_ =* 0.056 μm^2^/s (Fig. 1D and fig. S2). In contrast, all MEG-3 molecules exhibited restricted long-lived trajectories with an average apparent diffusion coefficient of *Dc =* 0.0018 μm^2^/s (Fig. 1C and D, and fig S2). In 3/3 cases where we captured the trajectories of three labeled MEG-3 molecules in the same P granule, their relative position remained fixed over time (Fig. 1C and D, and movie S2). Together these observations confirm that PGL-3 molecules exist primarily in a dynamic liquid-like phase, whereas MEG-3 molecules experience much slower dynamics resembling solid clusters within our experimental time scales.

Solid particulates that adsorb to liquid surfaces reduce surface tension and stabilize emulsions against coarsening (so-called “Pickering agents” (*30*)). To explore whether MEG-3 exhibits properties of a Pickering agent, we first tested whether MEG-3 affects the interfacial properties of PGL-3 condensates assembled *in vitro*. “Naked” PGL-3 droplets readily wet the surface of untreated glass slides (Fig. 1E and fig. S3A). In contrast, PGL-3 droplets coated with MEG-3 clusters exhibited lower wetting on glass slides, consistent with MEG-3 clusters adsorbing to the PGL-3 interface (Fig. 1E and fig. S3A). To determine whether MEG-3 stabilizes the PGL-3 emulsion against coarsening, we examined the evolution of a newly assembled PGL-3 emulsion overtime. We found that over the course of 180 minutes, the PGL-3 emulsion coarsens: droplets increased in size on average and decreased in number without a change in the total volume of PGL-3 in droplets (Fig. 1F and G, and fig. S3B to D). Addition of MEG-3 reduced coarsening, stabilizing droplet size and number over the 180 minutes of the experiment (Fig. 1F to H, and fig. S3B and C). Reduction in surface tension is correlated to the amount of surface occupied by the Pickering agent (*31*). As expected, we observed a correlation between MEG-3 concentration and the size of PGL-3 droplets in the stabilized emulsions (fig. S3E to I). We conclude that MEG-3 stabilizes the PGL-3 emulsion in a concentration-dependent manner, consistent with MEG-3 acting as a classic Pickering agent *in vitro*.

Even in the absence of MEG-3, coarsening of the PGL-3 emulsion was slow, occurring on a minute-to-hour time scale. Previous studies have shown that PGL-3 condensates are “aging Maxwell fluids” whose viscosity strongly increases overtime, eventually adopting glass-like properties (see Supplemental Text, (*2*, *32*)). To further explore the properties of the PGL-3 phase, we examined the dynamics of PGL-3 condensates challenged with excess PGL-3 protein. We found that within less than 1 minute of assembly, PGL-3 condensates became refractory to incorporation of new PGL-3, causing excess PGL-3 to form separate condensates (Fig. 2A). Mixing between “old” and “new” PGL-3 occurred progressively, indicating that PGL-3 molecules continue to exchange between the condensed and dilute phases (Fig. 2B). Addition of MEG-3 reduced coarsening but did not affect significantly the rate of old and new PGL-3 mixing, as expected for a surface agent that does not block exchange at the PGL-3 interface (Fig. 2A and C). These observations are consistent with PGL-3 condensates rapidly (within seconds) approaching kinetic arrest, where the soluble-to-condensate rate slows down significantly, limiting growth and coarsening.

**Fig. 2.**
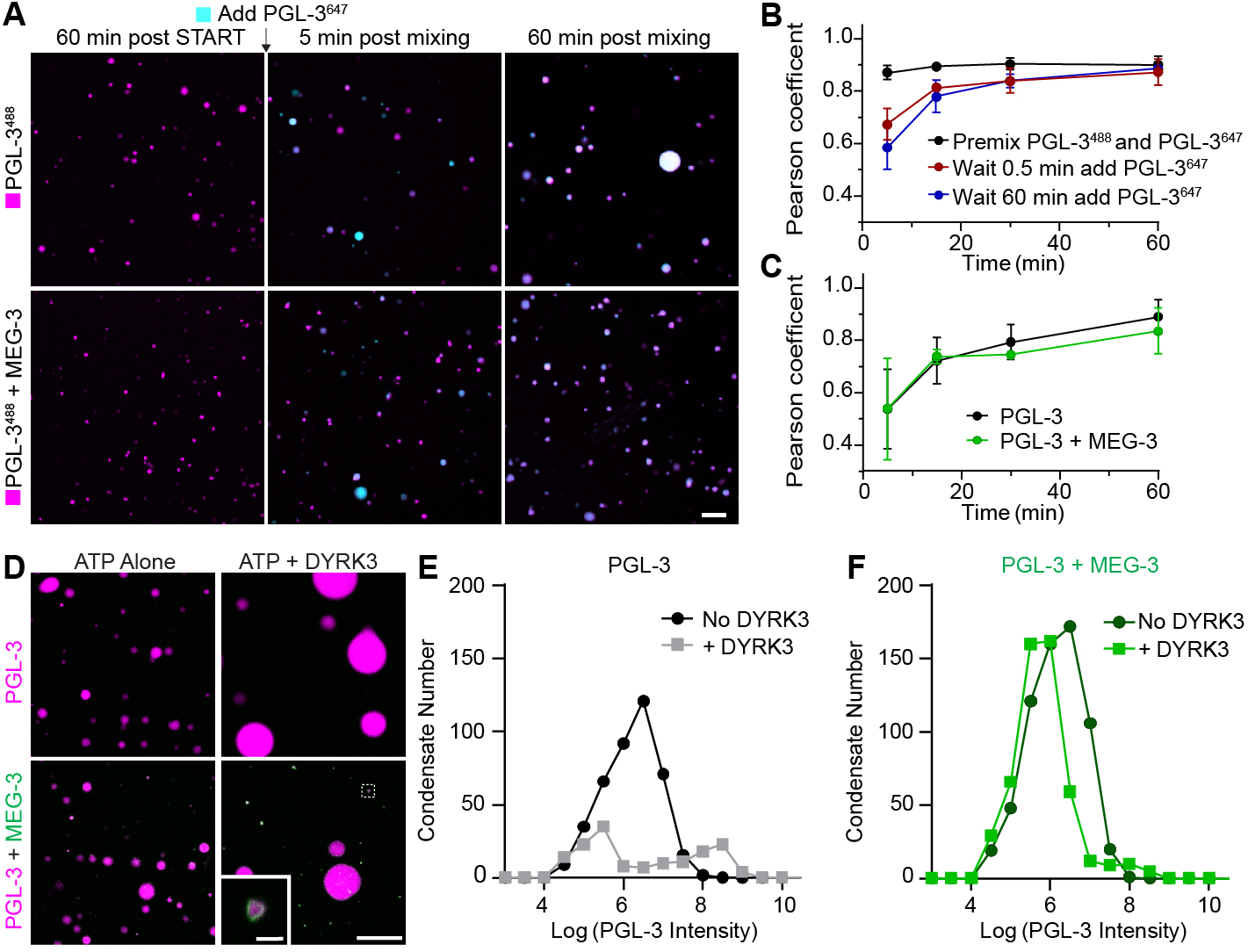
MEG-3 stabilizes PGL-3 condensates against kinase accelerated coarsening. A) Photomicrographs of a PGL-3^488^ emulsion (magenta) 60 min post initial assembly and at indicated time points after addition of new PGL-3^647^ (cyan). 2.5 μM PGL-3^488^ and 80 ng/μL *nos-2* RNA were incubated for 60 min in the presence or absence of 0.5 μM MEG-3 (not shown) before adding of 2.5 μM PGL-3^647^. Scale bar is 20 μm. B-C) Graphs showing the co-label correlation between PGL-3^488^ and PGL-3^647^ over time. (B) PGL-3^488^ emulsions were assembled as in (A) with zero, 0.5 min or 60 min of pre-incubation before addition of new PGL-3^647^. (C) PGL-3^488^ emulsions were assembled as in A (60 min pre-incubation) in the absence (black) or presence of MEG-3 (green). Each data point represents the Pearson coefficient averaged over 8 images. Error bars represent SD. D) Photomicrographs of PGL-3 (magenta) condensates assembled with and without 50 nM MEG-3 (green) and captured 60 min after addition of 100 μM ATP with and without 100 nM DYRK3. Inset shows a 7x magnification of the region shown in the dashed line box. Scale bars are 50 μm and 5 μm respectively. E) Histograms of PGL-3 condensates incubated for 60 min in 100 μM ATP with and without 100 nM DYRK3. Circles indicate the number of PGL- 3 condensates binned by the Log(Intensity) of each condensate captured from 20 images. Histograms of PGL-3 condensates assembled with 100 nM MEG-3 and incubated for 60 min in 100 μM ATP with 100 nM DYRK3. Circles indicate the number of PGL- 3 condensates binned by the Log(Intensity) of each condensate captured from 20 images. See fig. S4G and H for additional MEG-3 concentrations.

The slow soluble-to-condensate conversion rate of the PGL-3 emulsion is consistent with the apparent stability of P granules during most of germline development and suggests that active processes must operate to dissolve P granules during oocyte maturation and polarization. MBK-2 is required for P granule dissolution during polarization and its mammalian homolog DYRK3 has also been implicated in the dissolution of condensates in mammalian cells (*27*, *33*, *34*). We found that recombinant DYRK3 phosphorylates PGL-3 efficiently *in vitro* (fig. S4A). Addition of DYRK3 to pre-assembled PGL-3 condensates (2.5 μM) led to their complete dissolution over 60 minutes (fig. S4B to E). At 5 μM PGL-3, condensates could be maintained in the presence of DYRK3, but coarsened rapidly such that by 60 minutes only a few, large PGL-3 droplets (>5 μm) remained (Fig. 2D to E). Addition of MEG-3 did not affect the ability of DYRK3 to phosphorylate PGL-3 (fig. S4F), yet changed coarsening dynamics. In the presence of MEG- 3, after 60 minutes, numerous small PGL-3 droplets (<1 μm) could be detected in addition to large PGL-3 droplets (Fig. 2D and F, and fig. S4G and H). The small droplets were covered with MEG-3 and remained stable for an additional 60 minutes. These observations indicate that 1) phosphorylation enhances the soluble-to-condensate conversion rate of PGL-3 and accelerates coarsening and 2) MEG-3 is sufficient to stabilize small PGL-3 condensates even under conditions where PGL-3 dynamics (“fluidity”) have been increased by phosphorylation.

To examine the impact of MEG-3 on P granule dynamics *in vivo*, we used quantitative live-cell imaging to measure the number and size of PGL-3 condensates in wild-type and *meg-3 meg-4* mutants. We began by imaging eggs *in utero* as they progress through oocyte maturation and fertilization (Fig. 3A). We found that the volume of PGL-3 in condensates decreased 10-fold during oocyte maturation and remained low in newly fertilized zygotes as they complete the meiotic divisions (Fig. 3B to D and fig. S5A). Total PGL-3 levels did not change during this period (fig. S5B) consistent with a transient increase in PGL-3 solubility. We observed the same decrease in PGL-3 condensate volume in wild-type and *meg-3 meg-4* oocytes, indicating that the increase in PGL-3 solubility during the oocyte-to-zygote transition is not dependent on *meg-3 meg-4*. The size distribution of PGL-3 condensates post-dissolution, however, was different in the two genotypes. The PGL-3 emulsion coarsened rapidly in *meg-3 meg-4* zygotes with fewer larger condensates dominating the emulsion (Fig. 3D). In contrast, wild-type zygotes maintained many small PGL-3 condensates, consistent with MEG-3 stabilizing the PGL-3 emulsion against coarsening (Fig. 3D). Zygotes in late meiosis can survive outside of the uterus allowing for the acquisition of high-resolution images *ex utero*. These images confirmed that wild-type zygotes contain dozens of <1 μm condensates not observed in *meg-3 meg-4* zygotes (Fig. 3A and E).

**Fig. 3.**
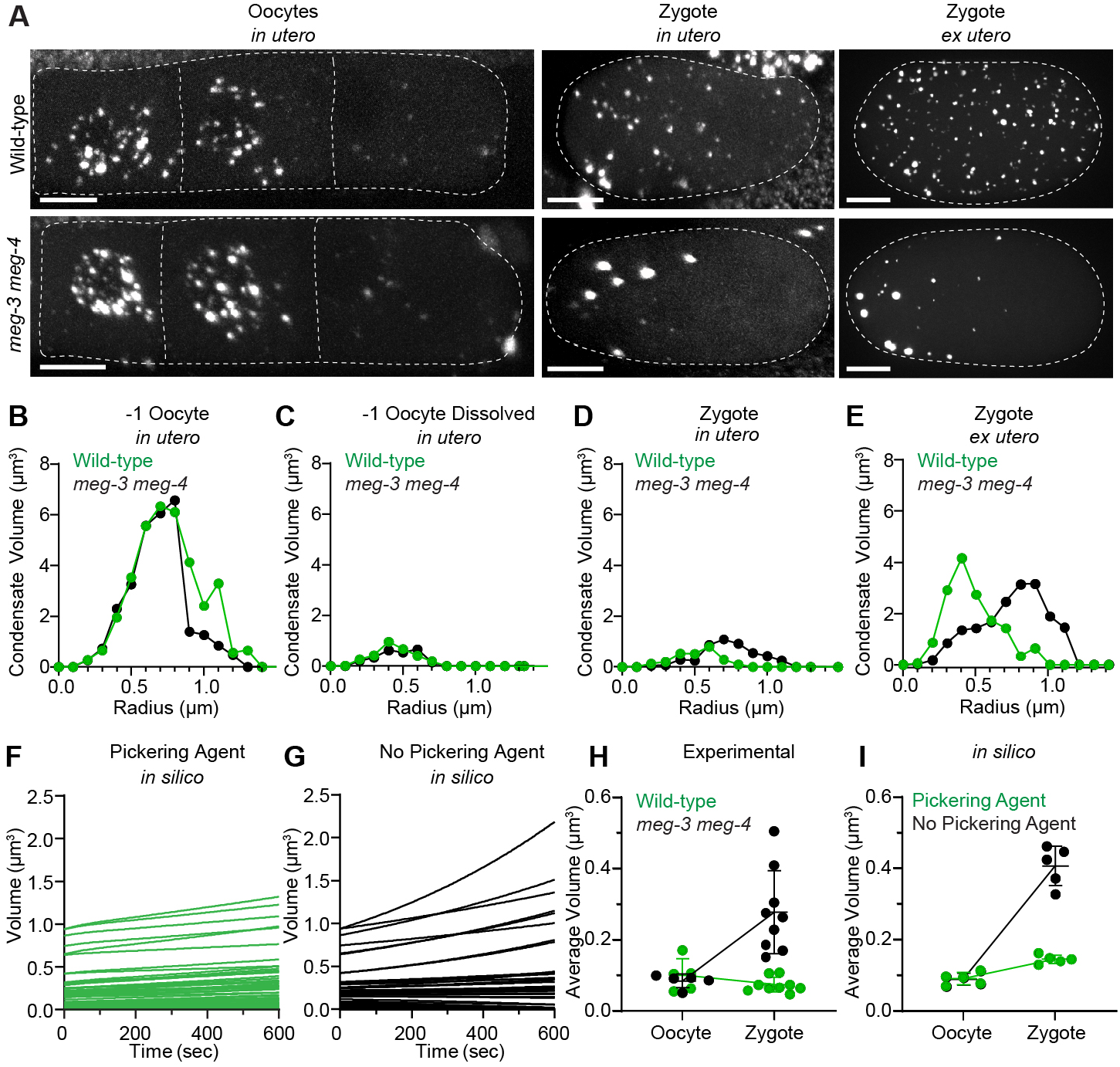
MEG-3/4 stabilize P granules against coarsening during the oocyte-to-zygote transition. A) Photomicrographs of wild-type and *meg-3 meg-4* oocytes and zygotes expressing PGL-3::mCherry (white). Photomicrographs were captured in live adult hermaphrodites (*in utero*) or after dissection out of the uterus (*ex utero*). Representative photomicrographs represent ~20% of oocyte volume and ~80% of zygote volume. The anterior (left) bias for PGL condensates in *meg-3 meg-4* zygotes correlates with anterior displacement of the oocyte nucleus (and associated P granules) that occurs immediately prior to fertilization. Scale bars are 10 μm. B-E) Histograms of PGL condensate volumes measured from images captured as in A representing 100% of oocyte and zygote volumes. Circles indicate the volume of individual PGL-3 in condensates binned by condensate radius in wild-type (green) and *meg-3 meg-4* (black) oocytes and zygotes. Volumes are higher in E compared to D due to higher detection sensitivity *ex utero*. F-G) Graphs showing the evolution of individual PGL condensates in a 10 min period starting after dissolution in wild-type and *meg-3 meg-*4 mutants under simulated conditions. Each line represents the evolution of the volume of a single condensate over time. H-I) Graphs showing the average volume of individual PGL condensates in oocytes and zygotes under experimental and simulated conditions. H) Each dot corresponds to an oocyte (same data set as shown in C) or zygote (same data set as shown in E) of the indicated genotypes. I) Each dot corresponds to one simulation. Simulations were run in the presence or absence of the Pickering agent.

These observations suggest that MEG-3 functions as a Pickering agent for the PGL-3 emulsion. To examine the physical plausibility of this hypothesis, we modeled *in silico* the kinetics of an idealized PGL-3 emulsion in the presence or absence of a Pickering agent that lowers surface tension. To account for the intrinsic tendency of PGL-3 towards a slow soluble-to-condensate conversion rate, we modeled PGL-3 dynamics under a “conversion-limited” scheme, where the soluble-to-condensate conversion rate is > 10-fold slower than the diffusion-limited adsorption/desorption rate of PGL-3, and therefore rate limiting for condensate growth/degrowth (see Supplemental Text). To model dissolution of PGL-3 condensates during oocyte maturation, we assigned a relatively high conversion rate to PGL-3 condensates and a high critical concentration for PGL-3 phase separation since most PGL-3 condensate dissolve during this period. To model MEG-3 as a Pickering agent, we reduced by a 100-fold the capillary length of MEG-3-coated PGL-3 condensates. We used the model to run simulations tracking the dynamics of condensates with starting sizes matching the distribution of PGL-3 condensates in oocytes post-dissolution (Fig. 3F and G). Coarsening was strikingly different in the presence or absence of the Pickering agent, with stabilization of smaller condensates requiring the Pickering agent. The simulations reproduced the increase in average condensate volume observed in *meg-3 meg-4* zygotes but not wild-type zygotes (Fig. 3H and I). These results support the hypothesis that MEG-3 stabilizes the PGL-3 emulsion against the increase in PGL-3 dynamics in zygotes by lowering the surface tension of PGL-3 condensates.

Next, we examined PGL-3 dynamics as zygotes transition from unpolarized to polarized. During this period, zygotes exit meiosis, assemble pronuclei that migrate to the center of the zygote, fuse and initiate the first mitotic division (Fig. 4A). During pronuclear formation and fusion, we observed new PGL-3 condensates appearing throughout the cytoplasm in both wild-type and *meg-3 meg-4* zygotes (Fig. 4B and C, and fig. S6A). The total volume of PGL-3 in condensates also increased (fig. S6B), without a change in total PGL-3 (fig. S6C), consistent with a return to low PGL-3 solubility in both genotypes. During this period, total condensate volume in the anterior half of the zygote decreased steadily eventually reaching zero, while total condensate volume in the posterior increased (Fig. 4D). Remarkably, in *meg-3 meg-4* zygotes, we also observed a decrease and increase in total condensate volume in the anterior and posterior, respectively, but the amplitude of the change was greatly reduced (Fig. 4E). By mitosis, in wild-type, all PGL-3 condensates were restricted to the posterior cytoplasm; in contrast, in *meg-3 meg-4* zygotes, PGL-3 condensates remained stable throughout the cytoplasm through the first cell division, suggesting that in *meg-3 meg-4* mutants, the PGL-3 emulsion is no longer responding efficiently to the MEX-5-driven solubility gradient (Fig. 4A to E).

**Fig. 4.**
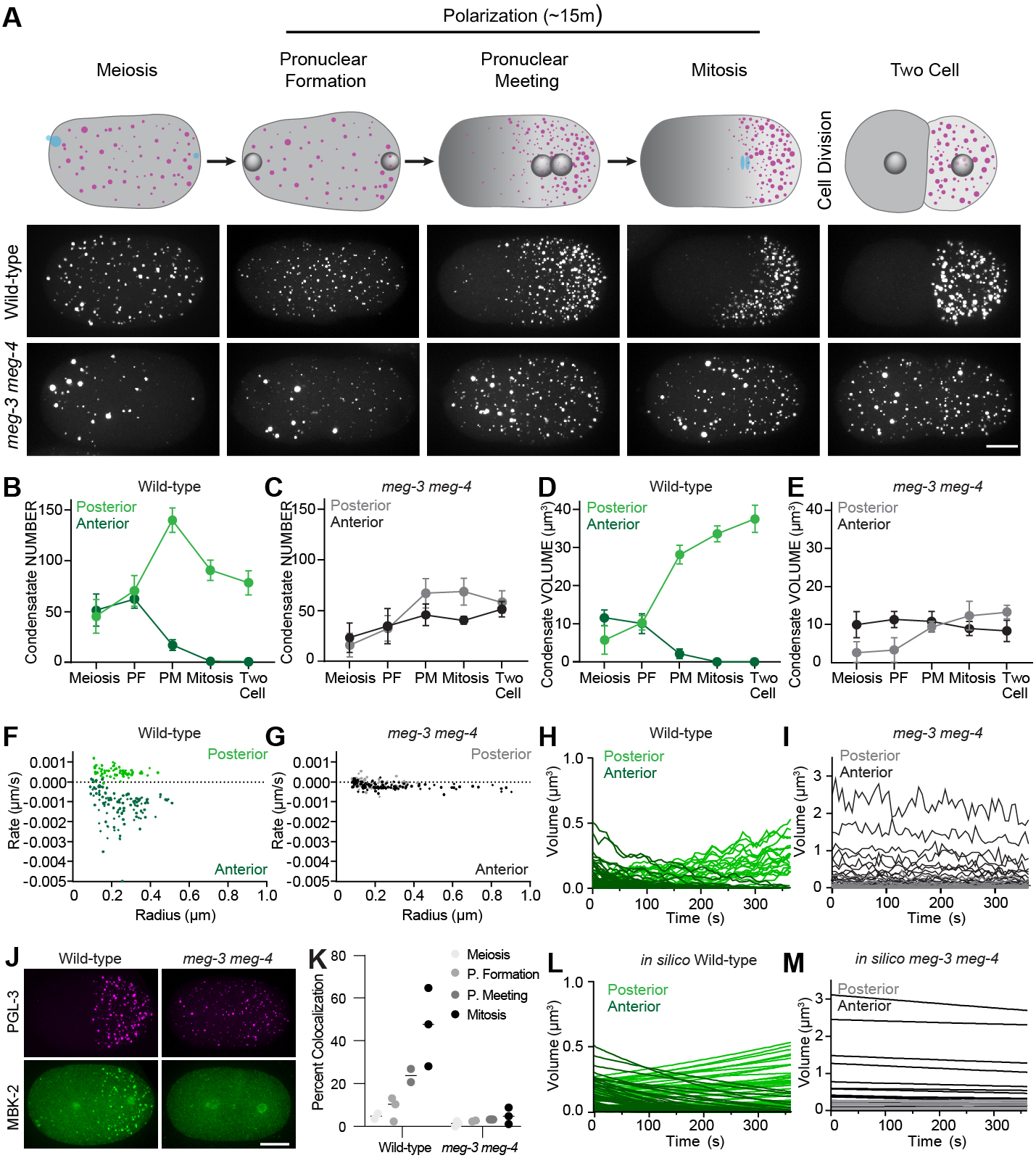
MEG-3/4 drive asymmetric growth of the P granule emulsion during polarization. A) Top row: Cartoons depicting PGL-3 condensates (red) and MEX-5 (grey) at different stages during the transition from unpolarized to polarized zygote. Bottom two rows: Photomicrographs of wild-type and *meg-3 meg-4* zygotes (*ex utero*) expressing PGL-3::mCherry and matching the stages shown in the cartoons above. Scale bar is 10 μm. B-E) Graphs showing the total number (B-C) and total volume (D-E) of PGL-3::mCherry condensates in the posterior (light color) or anterior (dark color) half of wild-type and *meg-3 meg-4* zygotes calculated from photomicrographs as in A. Circles represent the average from 5 zygotes. Error bars represent the SD. F-G) Graphs showing the rate of change in radius of PGL-3::mCherry condensates in wild-type and *meg-3 meg-4* zygotes calculated from traces as in H and I. H-I) Graphs showing the evolution of individual PGL-3::mCherry condensates in anterior and posterior during polarization in wild-type and *meg-3 meg-4* zygotes. Traces begin at pronuclear formation and end just prior to pronuclear meeting. J)Photomicrographs of fixed zygotes (mitosis) of indicated genotypes showing the distribution of MBK-2::OLLAS (green) and PGL-3::mCherry (magenta). Note that, in addition to P granules, MBK-2 localizes to centrosomes. K)Graph showing the percent of PGL-3:mCherry condensates colocalized with Ollas::MBK-2 puncta in wild-type or *meg-3 meg-4* zygotes at the indicated developmental stages. Each circle represents one zygote (>50 puncta) and each line represents the mean. L-M) Graphs showing the evolution of individual anterior (dark color) and posterior (light color) PGL-3 condensates under conditions simulating “wild-type” (starting condensate sizes as in wild-type zygotes, high conversion rate and Pickering agent) and “*meg-3 meg-4*” starting condensate sizes as in *meg-3 meg-4* zygotes, low conversion rate, no Pickering agent). Compare to experimental data in H and I.

To examine condensate dynamics directly, we tracked individual PGL-3 condensates in live zygotes from pronuclear formation to pronuclear meeting (Fig. 4F to I, and movies S3 and S4). These observations confirmed that, in *meg-3 meg-4* zygotes, condensates experience very slow growth/dissolution rates during polarization, with a slight bias for decay in the anterior (*k^A,avg^* −0.19 nm/s, *k^P,avg^* = 0.06 nm/s, Fig. 4G and I). Growth/decay rates were more than 5-fold faster in wild-type zygotes with a clear bias for dissolution in the anterior and condensation in the posterior (*k^A,avg^*-1.18 nm/s, *k^P,avg^* = 0.50 nm/s, Fig. 4F and H). These findings suggest that, unlike in oocytes where PGL-3 dissolution occurs independently of *meg-3* and *meg-4*, during polarization *meg-3* and *meg-4* are required to accelerate PGL-3 condensate dynamics.

In addition to *meg-3* and *meg-4*, P granule dissolution during polarization requires the DYRK kinase MBK-2 (*35*, *36*). Genetic epistasis experiments have shown that MBK-2’s dissolution activity is dependent on *meg-3* and *meg-4* (*27*) (Supplementary text). Consistent with these observations, using an epitope tagged allele of endogenous MBK-2, we found that MBK-2 is recruited to PGL-3 condensates during polarization and this recruitment is enhanced by *meg-3* and *meg-4* (Fig. 4J and K). MEG-3 and MBK-2 levels varied more than five-fold among PGL-3 condensates (fig. S7A to D). Condensate growth rates during polarization were also heterogeneous in a manner that did not correlate with initial condensate size (Fig. 4F and G). Together, these observations suggest that heterogeneous recruitment of MEG-3/MBK-2 variably fluidize PGL-3 condensates during polarization.

Based on these findings, we built on our theoretical model of the PGL-3 emulsion adding three new features specific to polarization: 1) variable soluble-to-condensate conversion rates for PGL-3 condensates to reflect heterogeneous fluidization by MEG-3/MBK-2, 2) higher solubility for PGL-3 in anterior cytoplasm to reflect the influence of the MEX-5 gradient, and 3) lower surface tension/capillary lengths for posterior condensates to reflect sustained coverage of posterior condensates by MEG-3 (“asymmetric Pickering agent”). Parameters were benchmarked to allow for complete dissolution of anterior granules in ~6 minutes as observed *in vivo*. We used the model to run simulations tracking the growth and decay of condensates matching the size distribution of PGL-3 condensates *in vivo* pre-polarization. The simulations faithfully recapitulated the coarsening-free dissolution of anterior condensates and growth of posterior condensates observed in wild-type zygotes (Fig. 4L and movie S5). Simulations mimicking conditions in *meg-3 meg-4* mutants reproduced the slow dynamics observed in those mutants (Fig. 4M and movie S6). The simulations also reproduced the rapid increase in PGL-3 condensate volume (fig. S8A) that is observed during mitosis in wild-type but not *meg-3 meg-4* zygotes (fig. S5A and S6B). The rapid rise is a consequence of the faster approach to equilibrium in the supersaturated environment of the posterior cytoplasm by PGL-3 condensates “fluidized” by MEG-3/MBK-2.

To test the importance of each feature in the model, we reran the wild-type simulations changing one model feature at a time. Simulations where condensates were assigned uniformly low conversion rates yielded decay rates that were too slow to clear anterior granules under the *in vivo* time constraints (fig. S8B and movie S7). Simulations with low PGL-3 solubility across the cytoplasm led to coarsening of anterior condensates (fig. S8C and movie S8). Simulations lacking the Pickering agent led to coarsening of posterior condensates (fig. S8D and movie S9). Together with the *in vitro* and *in vivo* findings, the theory supports a model where MEG-3 facilitates P granules polarization in response to the MEX-5-induced PGL-3 solubility gradient by accelerating PGL-3 conversion dynamics (via recruitment of MBK-2) and by functioning as a Pickering agent to lower surface tension on posterior condensates thus preventing coarsening during periods of high fluidity (see Supplemental Text for a full discussion of theoretical considerations).

Since their description by Ramsden and Pickering in the early 1900s (*18*, *19*), Pickering agents have been used widely to stabilize emulsions in the pharmaceutical, energy and food industries (*22*–*24*). Unlike surfactants (amphiphilic molecules that insert at interfaces), Pickering agents are nanoscale solid particulates that adsorb to interfaces upon partial wetting by both phases. Adsorption is energetically favored and balances the drive to reduce interfacial area, stabilizing the emulsion against Ostwald ripening and coalescence (*31*). Many types of solid particulates have been shown to function as Pickering agents, from silica to denatured proteins (*37*–*39*). The first described intrinsically disordered protein, casein, functions as a natural Pickering agent in homogenized milk (*40*). To our knowledge, the intrinsically disordered protein MEG-3 is the first characterized example of an *intracellular* Pickering agent and we speculate that other self-assembling biopolymers will exhibit similar properties. In somatic cells, PGL droplets are covered by EPG-2 clusters that may function like MEG-3 to regulate the size and dynamics of PGL droplets in preparation for autophagy (*41*). Artificial protein-RNA assemblies that adsorb to the surface of stress granules have been reported to influence their size and coalescence (*42*). mRNAs that accumulate on the surface of protein condensates could also serve as stabilizing agents (*43*, *44*). Given the rich diversity of biopolymers in cells, it is tempting to speculate that biopolymers acting as Pickering agents will prove a general organizing principle for biomolecular condensates. By interacting with enzymes like the DYRK kinase MBK-2, biological Pickering agents also regulate interfacial exchange to control the influx and efflux of molecules in and out of condensates in response to environmental changes. In this respect, surface-adsorbed biopolymers fulfill a boundary function for biomolecular condensates, reminiscent of the role of lipid bilayers and associated machineries (e.g. channels) in membrane-bound organelles. In fact, it has been speculated that membranes first arose from liposomes that functioned as Pickering stabilizers for aqueous emulsions (*45*).

## Acknowledgments

We thank the Johns Hopkins Integrated Imaging center (S10OD023548) for microscopy support. We thank the Lavis lab for HaloTag Ligands JF_549_ and JF_646_, the Griffin lab for MEG-3::Halo strain, We thank the Waugh Lav for TEV protease (pRK793, Addgene), Tony Hyman, the Baltimore Worm club and the Seydoux lab for many helpful discussions.

## Funding

This work was supported by the National Institutes of Health (GS: Grant number R37HD037047, AP: F32GM134630). GS is an investigator of the Howard Hughes Medical Institute.

## Author Contributions

A.W.F., A.P., and G.S. designed the research. A.W.F. and A.P. performed all experiments, collected, and analyzed data. C.L. performed the theoretical analysis. A.W.F., A.P., C.L., and G.S. prepared the manuscript with contributions from all authors.

## Competing interests

G.S. serves on the Scientific Advisory Board of Dewpoint Therapeutics, Inc.

## Data and materials availability

Source data is available in the manuscript or the supplementary materials. Other data supporting the findings of this study are available from the corresponding author upon reasonable request.

## Notes

### Competing Interest Statement

Geraldine Seydoux serves on the Scientific Advisory Board of Dewpoint Therapeutics

