## Supplementary Materials for "Pickering stabilization of a dynamic intracellular emulsion"

### Materials and Methods

#### *C. elegans* strains

*C. elegans* were cultured according to standard methods (Brenner, 1974). MEG-3, PGL-3, and MBK-2 were tagged at the endogenous locus with fluorescent or epitope tags using genome editing (*46*). Strains are listed in Table S1.

#### Laser scanning confocal microscopy

Super-resolution microscopy was performed using a Zeiss LSM 880 microscope equipped with a 63X-1.4 numerical aperture (NA) objective. Super-resolution images were acquired using ZEN software in SR mode, 1.8X zoom, and 0.15 μm Z-stack step size. Samples were illuminated with 488/561/637 nm solid-state laser, using a 488/561/633 transmitting dichroic (Zeiss) and a BP420-480+BP495-550 nm or P420-480+BP495-620 nm bandpass filter (Zeiss). The raw data was processed using default 3D Airyscan settings with ZEN software.

#### Spinning disk confocal microscopy

Super-resolution imaging of the PGL-3 emulsion *in vitro* and in eggs was performed using custom built inverted Zeiss Axio Observer with CSU-W1 SoRa spinning disk scan head (Yokogawa), 1X/2.8x/4x relay lens (Yokogawa), fast piezo z-drive (Applied Scientific Instrumentation), and an iXon Life 888 EMCCD camera (Andor). Samples were illuminated with 405/488/561/637 nm solid-state laser (Coherent), using a 405/488/561/640 transmitting dichroic (Semrock) and a 624-40/692-40/525-30/445-45 nm bandpass filter (Semrock). Images were taken using Slidebook software using a 40x-1.3NA/63X-1.4NA/100x-1.46NA objective (Zeiss) and a 1x, 2.8x, or 4x relay lens (Yokogawa) respectively depending on the sample. All spinning disk confocal imaging was performed in a climate controlled room kept at 18°C. See sections below for imaging analyses.

#### Near-TIRF single molecule microscopy

*HaloTag labeling*

HaloTag labeling was performed by feeding hermaphrodites expressing either MEG-3::Halo (EGD364) or PGL-3::Halo (JH3913) a concentrated OP50 culture (50μl) containing 0.25 μM of $JF_{549}$-HaloTag ligand and 1 μM $JF_{646}$-HaloTag ligand on NNGM plates (IPM Scientific) for at least 3 hrs at 20°C. Hermaphrodites were washed 3x in M9 solution and dissected to release zygotes for mounting under a #1.5 22-mm square coverslip (170 ± 5 μm, Marienfeld) in 5 μl of

M9 containing ~200 polystyrene bead (15.6 ± 0.03 μm diameter, Bangs labs). Two cell zygotes were chosen for analysis. Progression through the cell cycle is too fast in one cell zygotes to permit single molecule measurements.

*TIRF Imaging (Fig. 1C and D)*

Samples were simultaneously illuminated with 568 nm and 640 nm solid-state lasers and images were acquired in 'simultaneous' mode in the OMX Master software (GE Healthcare) on an DeltaVision OMX microscope with a Ring-TIRF module (GE Healthcare), 60x Olympus APO N TIRF/NA 1.49 oil objective and two front illuminated sCMOS cameras (PCO AG). For PGL-3::Halo (JH3913) time lapse images were acquired at an exposure of 100 ms or 150 ms at a time interval of 100 ms or 150 ms respectively. For MEG-3:Halo (EGD364) time lapse images were acquired at an exposure of 100 ms at a time interval of 500 ms or 1 s respectively. For all experiments, the ring diameter value, capture exposure setting, and time lapse frequency were chosen empirically to maximize signal intensity in each sample during the time course. (movies S1 and S2)

*Single Molecule Analysis (Fig. S2A)*

Single particle tracking of MEG-3::Halo (EGD364) and PGL-3::Halo (JH3913) was performed using the plugin TrackMate in Fiji (*47*). Feature size, and minimum intensity were empirically chosen so that >90% of the visible $JF_{549}$ labeled particles were detected. Trajectories that occurred outside the $JF_{646}$ labeled P granule area were manually excluded from analysis. 3D trajectory coordinates (x,y,t) were exported from Fiji. In some trajectories the $JF_{646}$ labeled P granules themselves moved slightly within the zygote overtime. To account for this drift, $JF_{549}$ particle trajectories were individually registered to the center of the $JF_{646}$ labeled P granule using Excel (Microsoft). Trajectory coordinates were loaded into Matlab (MathWorks) for further analyses. Mean squared displacement (MSD) curves were calculated for each track with at least 5 consecutive frames using MSDanalyzer (*48*). Individual MSD curves were fitted to the equation $MSD = 4D_c t^{\alpha}$ with an $R^2$ of at least 0.8.

**Protein and RNA purification**

MEG-3 full-length (aa1-862) fused to an N-terminal 6XHis tag in pET28a was expressed and purified from inclusion bodies as described (*49*). MBP::6XHis::TEV::PGL-3 was expressed and purified as described (*28*) with the following modification. MBP was cleaved using homemade TEV protease instead of commercial. A plasmid expressing 8XHis::TEV::8XArg tag protease was obtained from Addgene (PRK793) and purified as described (*50*). Before loading cleaved PGL-3 protein on to a heparin column (GE Healthcare), cleaved MBP::6XHis and 6XHis::TEV protease were removed using a HisTRAP column (GE Healthcare).

mRNAs were transcribed using T7 or SP6 mMessageMachine (Thermofisher) using manufacturer's recommendation as described (*49*). Template DNA for transcription reactions was obtained by PCR amplification from a plasmid containing the *nos-2* cDNA sequence (*49*). Free NTPs and protein were removed by lithium chloride precipitation. RNAs were resuspended in water and stored at −20°C. The integrity of RNA products was verified by agarose gel electrophoresis.

**Protein labeling**

Proteins were labeled with succinimidyl ester reactive fluorophores (Alexa Fluor™ 647 or DyLight™ 488 NHS Ester, Molecular Probes) following manufacturer instructions as described (*49*). Free fluorophore was eliminated by passage through three Zeba™ Spin Desalting Columns (7K MWCO, 0.5 mL) into protein storage buffer. The concentration of fluorophore-labeled protein was determined using fluorophore extinction coefficients measured on a Nanodrop ND-1000 spectrophotometer. Labeling reactions resulted in ~0.25-1 label per protein. Aliquots were snap frozen and stored. In condensation experiments, fluorophore-labeled protein was mixed with unlabeled protein for final reaction concentrations of ~12.5-50 nM of fluorophore labeled protein in the final reaction volume.

***in vitro* condensation experiments and analysis:**

Protein condensation was induced by diluting proteins out of storage buffer into condensation buffer containing 25 mM HEPES (pH 7.5), salt adjusted to a final concentration of 120 mM (30 mM KCl, 90 mM NaCl), and *nos-2* mRNA. Unless otherwise indicated, for co-assembly experiments 500 nM MEG-3, 3 µM PGL-3 and 80 ng/µL mRNA were used. PGL-3 and MEG-3 solutions contained 12.5-50 nM fluorescent trace labels with either Dylight™ 488, or Alexa Fluor™647 (indicated in figure legends).

PGL-3 wetting *in vitro* (Fig. 1E and fig. S3A).

Condensation reactions were incubated in a 1.5 mL eppendorf tube for 30 min at room temperature (18°C) before spotting 15 µL onto a No. 1.5 glass bottom dish (Mattek). Fast 3D image stacks (0.2 µm Z steps, ~5 µm Total) were captured every ~14 seconds using Slidebook software using a 63x oil objective and 1x relay lens. Time series Z stacks were exported from Slidebook software and further analysis was done in Imaris. Using the “Ortho” slicer functionality a 2D time series of images of the ZX plane (side view) for condensates wetting on the glass surface was generated. These images were exported from Imaris and the interfacial angle at 64s post wetting was measured using the “Angle Tool” functionality in Fiji. Angle values for each condensate were exported from Fiji and statistically analyzed and graphed using GraphPad Prism.

PGL-3 coarsening and mixing *in vitro* (Fig. 1F, Fig. 2A, and fig. S3E)

Condensation reactions were incubated in a 1.5 mL Eppendorf tube at room temperature (18°C) for indicated time points, before spotting 10 µL onto a No. 1.5 glass bottom dish (Mattek). For experiments where excess PGL-3 was added, 2.5 µM PGL-3 trace labeled with Alexa647 and 80 ng/µL mRNA with or without 500 nM MEG-3 were incubated for indicated times (0, 30s, 60 min) and then excess PGL-3 trace labeled with Dylight488 was added to the reaction (Final concentration 2.5 µM for a total of 5 µM PGL-3). 3D image stacks (5µm Z steps) were captured using Slidebook software using a 63x oil objective and 1x relay lens.

DYRK3/PGL-3 kinase assay (Fig. 2D and fig. S4A to C, F)

For kinase treatment final PGL-3 concentrations of 2.5 µM and 5 µM were utilized. Recombinant MEG-3 was added as indicated (50 nM, 150 nM, and 250 nM). Recombinant GST::DYRK3 kinase was purchased ThermoFisher (#PV3837). Protein condensation was induced by diluting PGL-3 and MEG-3 (when applicable) out of storage buffer into condensation buffer containing 25 mM HEPES (pH 7.5), salt adjusted to a final concentration of 120 mM (30 mM KCl, 90mM NaCl), and 1 mM DTT. No RNA was included. Condensation reactions were incubated in a 1.5 mL eppendorf tube at room temperature (18°C) for fifteen min. At the fifteen min time point a solution containing ATP or GTP (100 µM final), $MgCl_2$ (200 µM final), and DYRK3 (100 nM final) was added to the reaction, and further incubated at 30°C for sixty min. To examine the DYRK3 phosphorylation state of the PGL-3 samples, kinase treated samples were resuspended in 1x SDS sample buffer (minus EDTA) and resolved via Phos-tag SDS-PAGE (Wako Chemicals, Japan). Migration of Alexa labeled proteins within the gel was visualized using non-confocal laser scanner (Typhoon 7000, GE Healthcare) (fig. S4A and F). To analyze the impact of DYRK3 phosphorylation on PGL-3 solubility, kinase treated sample were centrifuged at 13,000 r.p.m for 10 min. The resulting supernatant and pellet were resuspended in 1X SDS sample buffer and resolved via SDS-PAGE (fig. S4C). Migration of Alexa labeled proteins within the gel was visualized using non-confocal laser scanner (Typhoon

7000, GE Healthcare). To examine the consequence of DYRK3 phosphorylation on PGL-3 condensation,15 µL of kinase reaction was spotted onto a No. 1.5 glass bottom dish (Mattek). At 2.5 µM PGL-3 condensates fully dissolved in the presence of kinase, while at 5 µM PGL-3 large, coarsened condensates remained 2 hours after addition of the kinase. 3D image stacks (5 µm Z steps) were captured using Slidebook software using a 40x oil objective and 2.8x relay lens. Images were exported from Slidebook software and further analysis was done in Fiji.

*in vitro* condensate image analysis and quantification (Fig. 1G and H, Fig. 2B, C, E and F, fig. S3B to D and F to I, and fig. S4D, E, G, and H)

Images were exported from Slidebook software and further analyses was done in Fiji. Images used for quantification were single planes taken from a 5 image Z-stack starting at the surface of the glass dish. To quantify condensates, a mask was created by thresholding images, smoothing once, filtering out objects of less than 4 pixels to minimize noise, applying a watershed filter to improve separation of objects close in proximity, and converting to a binary image by the Otsu method using the nucleus counter cookbook plugin. Minimum thresholds were set to the mean intensity of the background signal of the image plus 1-2 standard deviations. The maximum threshold was calculated by adding 4-6 times the standard deviation of the background. Using generated masks, the 2D area $\mu m^2$ and the intensity of each condensate were measured and exported into Excel (Microsoft) for further analysis. Assuming that each condensate is a sphere in solution the radius (r) was determined from the 2D area $\mu m^2$ (A) measurement, $r = (A/\pi)^{0.5}$. This radius value was used to calculate 3D volume (v) and surface area (s.a.) for each condensate, $v = (4/3)\pi r^3$ and $s.a. = 4\pi r^2$. Objects from multiple images were identified and quantified as described above. Histograms of PGL-3 condensate volumes were generated by summing the volume binned by the radius of each PGL-3 object, and normalized to the total volume of condensates in the reaction. Final data points are the average of 4 replicates (a total of 20 images per replicate, Fig. 1G and H, Fig. S3B to D and F to I). To quantify the mixing of PGL-3 condensates over time, the Pearson's correlation coefficient of PGL-3 intensities in the 488 or 647 channels was compared. Final data points are the average of 3 replicates (40 images per replicate, Fig. 2B and C). For kinase assays, addition of the kinase resulted in the formation of large droplets that rapidly wet the surface of the coverglass resulting in inaccurate volume measurements. To compare the amount and distribution of condensates, histograms of PGL-3 condensate intensity were generated by taking the number of condensates binned by the intensity of each PGL-3 object. Final data points are the average of 4 replicates (a total of 5 images per replicate, Fig. 2E and F and fig. S4D, E, G, and H).

#### Live imaging experiments and analysis

Since the PGL-3 emulsion is temperature sensitive, live imaging was performed at 18°C, a temperature that preserves the integrity of PGL-3 condensates in wild-type (JH3914) and *meg-3 meg-4* (JH3915) zygotes (fig. S9A and B). Oocyte maturation and fertilization cannot occur *ex vivo* and therefore must be analyzed in hermaphrodites (*in utero*). Zygotes that have completed meiosis can develop outside of hermaphrodites, and therefore were analyzed after dissection (*ex utero*) to maximize the sensitivity of fluorescence imaging.

*in utero* analyses (Fig. 3A to D, and fig. S5A and B).

Adult hermaphrodites (JH3914, JH3915) were washed 2x in L-15 (Gibco) solution and were immobilized on a 35mm, No. 1.5 glass bottom dish (Mattek) in a L-15 medium containing 20 µm polystyrene beads (Polysciences) and 1 mM levamisole (Sigma). 3D image stacks spanning the entire germline and uterus (0.2 µm Z steps) were taken at max laser intensity using Slidebook software using a 40x oil objective and 2.8x relay lens. Images were exported from Slidebook and imported into Imaris image analysis software for further analysis. Individual oocytes and zygotes were isolated using the “Surface” functionality. The “Spot” functionality was used to identify P granules and statistically code the 3D sphere volume ($\mu m^3$) of each identified P granule within a zygote. Volume values for each zygote were exported from Imaris and statistically analyzed and graphed using GraphPad Prism. To calculate the total intensity of PGL-3 in oocyte and zygotes Imaris generated “Surfaces” were exported to Fiji. Images were background corrected and the total intensity was summed. The total intensity of each sample was then normalized to the average intensity of the sample.

*ex utero analyses* (Fig. 3A and E, Fig. 4B to E, and fig. S6A to C):

Adult hermaphrodites expressing endogenous PGL-3::mCherry (JH3914, JH3915) were washed 2x in L-15 (Gibco) solution and dissected to release the zygotes into ~100 µl L-15 (Gibco) solution on a 35mm, No. 1.5 glass bottom dish (Mattek) image stacks (0.2 µm Z steps, 10 µm total) were taken at max laser power at indicated developmental time points using Slidebook software using a 40x oil objective and 4x relay lens. Images were exported from Slidebook and imported into Imaris image analysis software for further analysis. In Imaris the “Spot” functionality was used to identify all P granules within the zygote and statistically code the 3D sphere volume (μm$^3$) of each identified P granule. Individual condensate volume values for P granules for each zygote were exported from Imaris and statistically analyzed and graphed using GraphPad Prism (Fig. 3E, fig. S6A and B). To calculate the total intensity of PGL-3 images were exported to Fiji. Images were background corrected and the total intensity was summed (fig. S6C). To quantify the number of condensates in the anterior of posterior of the zygote (JH3918 and JH3919), images were exported to Fiji (Fig. 4B to E). Using a maximum projection, a mask was created to identify PGL-3 condensates by smoothing, thresholding, filtering out objects of less than 2 pixels, applying a watershed filter to improve separation of objects close in proximity, and converting to a binary image by the Otsu method using the nucleus counter cookbook plugin. Using generated masks, the number and the area of each condensate was calculated. Assuming that each condensate is a sphere in solution the radius (r) was determined from the 2D area μm$^2$ (A) measurement, $r = (A/\pi)^{0.5}$. This radius value was used to calculate 3D volume (v) and surface area (s.a.) for each condensate, $v = (4/3)\pi r^3$ and $s.a. = 4\pi r^2$ . Condensates were split into either anterior or posterior half of the zygote, and plotted by stage.

*Live tracking* (Fig. 4F to I, and movies S3 and S4):

Live tracking of P granules in oocytes was not possible due to high photosensitivity and movement of the eggs from the oviduct to the spermatheca and into the uterus. Live tracking in zygotes (JH3914, JH3915) was performed *ex utero* by acquiring 3D image stacks (0.6 μm Z steps, 12 μm total) every ~5 seconds using Slidebook software using a 100x/1.46NA oil objective and 1x relay lens. Time series images were exported from Slidebook software. Images were corrected for photobleaching using the “Bleach Correction” feature in Fiji and further analyses were done in Imaris. Individual P granules were identified using the “Spot” functionality in Imaris. Spots were tracked over time using the Imaris “Autoregressive Motion” algorithm. Tracks were manually inspected and corrected if required. Volume and Z-coordinate values were exported from Imaris for further analysis in excel and graphed using GraphPad Prism. All tracks that were less than 5 frames, entered or exited the Z-plane during imaging, or moved from the posterior to the anterior (or the reverse) during imaging were discarded. To calculate the rate of change in size, radius was calculated from the volume for each P granule. The slope of the change in radius over time was calculated and plotted using GraphPad Prism.

**Immunostaining and analysis:**

For immunostaining of MEG-3::GFP and PGL-3::mCherry (Fig. 1B and fig. S1), gravid adult hermaphrodites (JH3921) were placed on #1.5 22x22 mm coverslip (170 ± 5 μm, Marienfeld) coated with 0.1% poly-L-lysine (Sigma) in a L-15 medium containing 25 μm polystyrene beads (Polysciences). The zygotes were extruded by squashing hermaphrodites with an opposing #2 24x50 mm glass coverslip (VWR) and snap frozen in liquid nitrogen. The 24x50 mm coverslip was removed and zygotes were permeabilized by incubation in chilled methanol at -20°C overnight. Coverslip were then rehydrated in PBS-Tween (0.1%) BSA (0.5%) (PBST/BSA) at room temperature for at least 3 hours. Coverslip were mounted onto a 25x75x1 mm microscope slide (ThermoFisher Scientific) in ~5 μl of antifade mounting medium (VectaShield). 3D image stacks (0.15 μm Z steps) were taken at indicated developmental time points using ZEN software using a 63x oil objective (Zeiss).

For immunostaining of OLLAS::MBK-2 and PGL-3::mCherry (Fig. 4J and K, and fig. S7C and D), gravid adult hermaphrodites (JH3916, JH3917) were placed on #1.5 22x22 mm coverslip (170 ± 5 μm, Marienfeld) coated with 0.1% poly-L-lysine (Sigma) in a L-15 medium containing 25 μm polystyrene beads (Polysciences). The zygotes were extruded by squashing hermaphrodites with an opposing #2 24x50 mm glass coverslip (VWR). These zygotes were snap frozen in liquid nitrogen. The 24x50 mm coverslip was removed and zygotes were permeabilized by incubation in chilled methanol at -20°C overnight. Coverslip were then blocked in PBS-Tween (0.1%) BSA (0.5%) (PBST/BSA) at room temperature for at least 3 hours, and incubated with Rat αOLLAS-L2 (1:200, Novus Biological Littleton, CO) at 4°C overnight. Following primary incubation, the coverslips were extensively washed with TBST. Secondary antibodies anti-Rat Alexa Fluor 488 (LifeTech) were diluted in PBST/BSA applied to coverslips at room temperature. Following secondary incubation, the coverslips were extensively washed with PBST. Coverslip were mounted onto a 25x75x1 mm microscope slide (ThermoFisher Scientific) in ~5 μl of antifade mounting medium (VectaShield). 3D image stacks (0.2 μm Z steps) were taken at indicated developmental time points using Slidebook software using a 40x oil objective and 4x relay lens. Images were exported from Slidebook software and further analyses was done in Imaris. In Imaris the "Spot" functionality was used to identify all MBK-2 and PGL-3 puncta within the zygote and statistically code the 3D sphere volume ($\mu m^3$). Using the 'measure' functionality, a nearest neighbor measurement from the center of these statistically coded MBK-2 and PGL-3 condensates was performed. PGL-3 and MBK-2 condensates that were within 500nm were considered colocalized, and counted. These values were exported from Imaris and statistically analyzed and graphed using GraphPad Prism.

Percent colocalization = (Total # colocalized PGL-3&MBK-2 condensates) / (Total # PGL-3 condensates) X 100.

For immunostaining of endogenous MEG-3::OLLAS and untagged PGL-3 (fig. S7A and B), gravid adult hermaphrodites (JH3477) were placed on a custom-made chambered slide (ThermoFisher) coated with 0.1% poly-L-lysine (Sigma), and zygotes were extruded by squashing with a glass coverslip. Zygotes were frozen on pre-chilled aluminum blocks for 5 min. The coverslip was removed, and zygotes were permeabilized by incubation in chilled methanol at -20°C for 15 min followed by incubation in chilled acetone -20°C for 10 min. Slides were then blocked in PBS-Tween (0.1%) BSA (0.5%) (PBST/BSA) at room temperature and incubated with primary antibodies at 4°C. Antibodies were diluted in PBST/BSA as follows: KT3 (1:10, DSHB), Rat αOLLAS-L2 (1:200, Novus Biological Littleton, CO). Primary antibodies were applied sequentially (OLLAS before KT3) to avoid cross reaction. Secondary antibodies Anti-Mouse DyLight 650 (Abcam) and anti-Rat Alexa Fluor 488 (LifeTech) were diluted in PBST/BSA applied to slides at room temperature. Zygote were mounted under a #1.5 22-mm square coverslip (170 ± 5 μm, Marienfeld) in ~5 μl of antifade mounting medium (VectaShield). 3D image stacks (0.2 μm Z steps) were taken at indicated developmental time points using Slidebook software using a 40x oil objective and 4x relay lens.

To quantify the ratio of MEG-3 or MBK-2 to PGL-3 signal in the zygote (fig. S7A to D), images were exported to Fiji. For MEG-3/PGL-3 images, maximum projections were analyzed. For MBK-2/PGL-3 images, single planes spaced 1.6 μm apart were analyzed because the cytoplasmic background signal of MBK-2 was high relative to the intensity of MBK-2 in condensates. For both data sets a mask was created to identify PGL-3 condensates by smoothing, thresholding, filtering out objects of less than 2 pixels, applying a watershed filter to improve separation of objects close in proximity, and converting to a binary image by the Otsu method using the nucleus counter cookbook plugin. Using generated masks, the integrated intensity within each object for both PGL-3 and MBK-2 or MEG-3 channels was calculated. To remove non-specific background signal the mean intensity of an image field calculated from a region outside of the zygote was subtracted from each pixel. The intensity of each condensate was divided by the intensity of the zygote to normalize to different staining efficiencies. The ratio of MBK-2 or MEG-3 to PGL-3 was calculated for each condensate and plotted versus the position on the anterior/posterior axis for multiple zygotes at each stage.

#### *in silico* modeling (Fig. 3F,G, and I, Fig. 4L and M, and fig. S8A to D)

For the oocyte to zygote simulation, 3 sets of starting condensate radii were generated from experimentally observed data (Fig. 3C). Using the mathematical formalism described in the Supplemental Text, the change in condensate radii was modeled over a 10 min window using a custom MatLab (MathWorks) program. This time window corresponds to fertilization of the oocyte and passage through the spermatheca into the uterus of the animal. To graphically display the simulation results, the radii output generated in MatLab (MathWorks) of individual condensates were converted to spherical volumes using Excel and plotted over time using GraphPad Prism. To display the size distributions of the simulated condensates, histograms of condensate volume were generated by summing the volume binned by the radius of each condensate at the start and end of the simulation.

For the polarization simulation, a representative starting P granule size distribution was generated from experimentally observed data (Fig. 4A) for both wild-type and *meg-3 meg-4* genotypes. The simulations shown in (Fig. 4L and fig. S8B to D) used a wild-type starting radii distribution, and simulation shown in (Fig. 4M) utilized a *meg-3 meg-4* starting radii distribution. Using the mathematical formalism described in the Supplemental Text, the change in condensate radii was modeled over a 6 min window using a custom MatLab (MathWorks) program. This time window corresponds to time frame encompassing the initiation of polarization at pronuclear formation until the zygote is polarized at pronuclear meeting. To plot the simulation results, output radii were converted to spherical volumes using Excel and plotted over time using GraphPad Prism. To visualize simulations results, output radii were converted to spherical volumes and randomly distributed in the anterior or posterior of an idealized zygote and visualized over time. Text and animation were added to these time series using Premiere Pro, movies S5 to S9.

**Analysis software**

Excel (Microsoft) and GraphPad Prism v9 were used for all data analysis and graphing. Images were captured with Slidebook v6 (Intelligent Imaging Innovations) or Zen Black (Zeiss). Fiji v1.52g, Imaris v9.51 (Bitplane), Matlab v2017b (Mathworks), and Zen Blue (Zeiss) were used for image processing and analysis. Matlab v2019b (MathWorks) was used for all *in silico* simulations. Premiere Pro v14.8 (Adobe) was used for video editing.

### Supplementary Text

#### Introduction to emulsions

Emulsions are collections of liquid condensates that co-exist in a common medium. We describe how liquid emulsions evolve over time.

***Diffusion-limited emulsions:***

Consider a simple polymer with several low-affinity binding sites (fig. S10A). The binding sites allow polymer molecules to bind to each other reversibly to form extended dynamic networks of weakly interacting molecules. At low concentration, polymer molecules exist primarily as dispersed soluble monomers. Above a certain concentration (“critical concentration”), polymer molecules phase separate and redistribute into dense condensates surrounded by a more dilute phase (*1*). Molecules in the condensates are bound dynamically to each other in a network that is constantly remodeling. Molecules in the dilute phase are free to diffuse and, when near the condensate surface, interact with molecules in the condensates. The **conversion rate** refers to the rate at which new molecules are incorporated into condensates from the dilute phase.

In highly dynamic emulsions, the conversion rate is faster than the rate of diffusion-limited adsorption of molecules in the dilute phase to the surface of the condensate (**“diffusion-limited”**). In a diffusion-limited system coarsening and growth occur rapidly, and the entire volume of the condensate exchanges rapidly with the dilute phase (fig. S10A).

***Conversion-limited emulsions:***

Recently, theoretical and experimental observations have indicated that certain polymer condensates evolve into less dynamic states overtime (*2*, *51*, *52*). As binding interactions increase between polymers in the condensates, the polymers become more and more intertwined with each other and less free to engage in binding interactions with molecules in the dilute phase (*2*, *8*, *53*). As the binding sites saturate, the conversion rate falls below the diffusion dependent adsorption rate of new monomers to the condensate surface. This makes it more difficult for molecules to enter the condensates (*2*, *8*). Under such **“conversion-limited”** scheme, exchange still occurs between the condensate and dilute phases, but net incorporation of new molecules into condensates slows down (fig. S10B).

***Coarsening:***

Molecules at the condensate surface make fewer favorable binding interactions with their neighbors and thus are energetically less stable than molecules deep inside the condensates. The energy associated with this difference is called surface tension. To minimize surface tension, emulsions tend to minimize overall interfacial area by favoring the growth of larger condensates (which have low surface-to-volume ratios) over that of smaller condensates (which have high surface-to-volume ratios). This thermodynamic evolution is called **coarsening**.

Two mechanisms drive coarsening: 1) coalescence (fusion) of condensates and 2) Ostwald ripening. Ostwald ripening refers to the preferential exchange of molecules from small to large droplets. Overtime small condensates are absorbed into large condensates until eventually the emulsion reduces to one large condensate (fig. S10C) (*54*).

The rate of coarsening depends on the surface tension, the solubility and diffusion rate of the polymer in the dilute phase, and the conversion rate. Coarsening driven by coalescence additionally requires interaction between condensates and depends on the density and diffusion of condensates. Increase in any of these parameters will increase the rate of coarsening and lead to fewer, larger condensates in a shorter time. Emulsions under the diffusion-limited coarsening scheme with high conversion and diffusion rates coarsen rapidly. In contrast, conversion-limited emulsions with slow conversion rates will typically coarsen over longer time scales, allowing emulsions to remain stable for hours or longer.

***Regulation of coarsening in cells:***

Several mechanisms have been proposed to regulate coarsening in cells. Reversible modifications (ie. phosphorylation) that transiently weaken binding interactions will increase the number of molecules in the condensate available for binding and thus increase the conversion rate (*55*). Modifications can also increase the solubility of polymer molecules in the dilute phase, thus increasing shedding of molecules by smaller condensates and Ostwald ripening (*54*).

Cellular structures that keep condensates away from each other prevent coalescence and slow down coarsening way (*5*, *13*, *56*). For example, an F-actin network stabilizes an emulsion of hundreds of nucleoli in *Xenopus laevis* oocytes, but when actin is disrupted, the emulsion rapidly coarsens (*5*). Coarsening has also been proposed to be reduced in the presence of cytoplasmic gradients that counteract Ostwald ripening by controlling diffusion in the dilute phase (*14–17*, *57*). Active mechanisms that continuously regenerate small condensates will also counteract coarsening (*14*).

In this study, we identify a new mechanism for coarsening regulation in cells by showing that the intrinsically-disordered protein MEG-3 functions as a Pickering agent for P granules, liquid condensates in *C. elegans* zygotes. Pickering agents were first described in the context of inorganic emulsions by Ramsey and Pickering (*18*, *19*). Working with oil in water emulsions, Ramsey and Pickering discovered that small inert particulates such as clay greatly slow down coarsening. **"Pickering agents"** are particulates ranging from 0.1 - 100 µm in diameter and are typically at least an order of magnitude smaller than droplets in the emulsion (*37*). They can be wetted by both the condensed and dilute phases leading to strong interfacial adsorption (*37*). The adsorption energy creates an energetically favorable counterbalance to the surface tension cost, effectively neutralizing the primary drive for coarsening (fig. S10C).

### Theoretical model

#### *Model assumptions*

The key model assumptions are:

1. PGL-3 condensates coarsen under the conversion-limited scheme (fig. S10D). We ignore coarsening by coalescence and fusion, as these events are infrequently observed experimentally *in vivo*.
2. MEG-3 clusters localizing to the surface of PGL-3 condensate reduce surface tension (Pickering effect).
3. PGL-3 condensate conversion dynamics are accelerated by unknown factors during the oocyte to zygote transition and by MEG-3-dependent recruitment of MBK-2 to PGL-3 condensates during polarization.
4. During polarization, PGL-3 condensates experience i) spatially differential surface tensions due to enrichment of MEG-3 in the posterior and ii) spatially differential levels of supersaturation due to enrichment of MEX-5 in the anterior.

#### *Justifications for model assumptions*

*Assumption 1:*
*PGL-3 condensates coarsen under the conversion-limited scheme* (fig. S10D). *We ignore coarsening by coalescence and fusion, as these events are infrequently observed experimentally in vivo.*

Standard diffusion-limited Ostwald coarsening kinetics (with reasonable parameters from the literature shown in Table S2) predicts a coarsening kinetics much faster than experimental observations. Therefore, we assume that condensate coarsening kinetics is conversion-limited (fig. S10D). This assumption is consistent with recent simulation work showing that biomolecular condensates are likely to be in slow kinetics (e.g., glassy) states (*58*, *59*). Specifically, the standard kinetics equations in the diffusion-limited regime are given by (*60*, *61*)

$$\frac{dR_i(t)}{dt} = \frac{A}{R_i}\left(\epsilon(t) - \frac{l_c}{R_i(t)}\right), \tag{1}$$

where $R_i(t)$ denotes the radius of the $i$-th condensate in the system, $l_c$ is the capillary length, $A$ is a constant quantifying the kinetic rate, and $\epsilon(t)$ is the dimensionless supersaturation level given by:

$$\epsilon(t) \equiv \frac{c(t)}{c_{\text{out}}} - 1\,, \tag{2}$$

with $c(t)$ being the far-field concentration in the dilute phase and $c_{\text{out}}$ is the equilibrium concentration in the dilute phase right outside a flat interface. On the other hand, the kinetics equations in the conversion-limited regime we consider here are of the form:

$$\frac{dR_i(t)}{dt} = K\left(\epsilon(t) - \frac{l_c}{R_i(t)}\right). \tag{3}$$

where $K$ is a kinetic rate that is independent to $R_i$ (unlike the standard formulation) due to the fact that the growth/shrinkage of condensates is not diffusion-limited.

A physical mechanism that can explain the slow internal dynamics in the condensates is the rugged energy landscape (*62*) caused by the binding and unbinding of protein domains within the condensates (fig. S10E). The binding and unbinding can be either due to thermal fluctuations (slow), or a phosphorylation/de-phosphorylation cycle (fast), leading to a kinase-controlled kinetic rates of condensate dynamics.

*Assumption 2*

*MEG-3 clusters localizing to the surface of PGL-3 condensates reduce surface tension (Pickering effect).*

This is supported by the experimental observations that MEG-3 clusters coat the surface of PGL-3 condensates, and that the surface tension of the resulting condensates is greatly modified *in vitro*.

*Assumption 3*
*PGL-3 condensate conversion dynamics are accelerated by unknown factors during the oocyte to zygote transition and by MEG-3-dependent recruitment of MBK-2 to PGL-3 condensates during polarization.*

This assumption is supported by examination of P granule dynamics in wild-type and *meg-3 meg-4* mutants (Fig. 4A to E) and additional genetic data presented in Wang et al. (*27*) (summarized below).

Oocyte to zygote transition: PGL-3 condensates dissolve in maturing oocytes, and this dissolution is observed in both wild-type and *meg-3 meg-4* mutants (Fig. 3A to C). These observations indicate that accelerated PGL-3 dynamics at this stage do not require *meg-3* or *meg-4*. Previous genetic studies have also shown that the DYRK kinase MBK-2 is not required for P granule dissolution at this stage (*27*). The factors that induce P granule dissolution at this stage are not known.

Polarization: In polarizing zygotes (pronuclear formation to pronuclear meeting), most P granules in the anterior cytoplasm dissolve and many P granules in posterior cytoplasm grow. These dynamics are greatly reduced in *meg-3 meg-4* mutants (Fig. 4). Prior genetic studies have shown that P granule dynamics at this stage also require the DYRK kinase MBK-2 (*27*). In zygotes lacking MBK-2, P granules do not dissolve in the anterior cytoplasm. MBK-2 is opposed by the phosphatase PP2A$^{pptr-½}$ . In zygotes lacking PP2A$^{pptr-½}$ , the opposite phenotype is seen: P granules dissolve throughout the cytoplasm of polarizing zygotes. Strikingly, the “hyper-dissolution” phenotype of zygotes lacking PP2A$^{pptr-½}$ can be suppressed by reducing either *mbk-2* or *meg-3* (*27*). These genetic results indicate that MEG-3/MEG-4 are required for MBK-2 activity on P granules. These findings are consistent with our observation that MEG-3 and MEG-4 are required to recruit high levels of MBK-2 to P granules during polarization (Fig. 4J and K).

*Assumption 4*

*During polarization, PGL-3 condensates experience i) spatially differential surface tensions due to enrichment of MEG-3 in the posterior and ii) spatially differential levels of supersaturation due to enrichment of MEX-5 in the anterior.*

Assumption 4i) follows from Assumption 2 and the observation that MEG-3 becomes progressively depleted from anterior PGL-3 condensates during polarization (Fig. S7A and B).

Assumption 4ii) follows from known effects of MEX-5 (*63*, *64*) which has been shown to increase the critical concentration for PGL-3 condensation in anterior cytoplasm, possibly by competing for RNA (which enhances PGL-3 condensation *in vitro*). This assumption is also supported by simulation results that demonstrate that, in the absence of a lower supersaturation state for anterior condensates, large PGL-3 condensates in the anterior grow transiently due to Ostwald ripening (fig. S8C), a behavior not observed in zygotes (Fig. 4H).

***Modeling Oocyte (O) to Zygote (E) transition***

<u>*Wild-type*</u>

Here, we focus on the last 10 min of the O to E transition, which marks the initiation of coating of MEG-3 onto PGL-3 condensates. Since the MEX-5 gradient is not yet present and MEG-3 is uniformly distributed, all condensates are treated as equal and the kinetic equations are given by (3), except to model non-instantaneous adsorption of MEG-3 clusters onto the PGL-3 surface, we assume that the capillary length decreasing from $l_c(t=0)=15$ nm to $l_c(t=600\ \text{sec})=0.15$ nm according to the following equation:

$$l_c(t) = (15\text{nm} - 0.15\text{nm})e^{-\frac{t}{\tau}} + 0.15\text{nm} \qquad (4)$$

where $\tau = 30$ sec. The small value of the capillary length is due to Pickering effects of MEG-3 clusters on PGL-3 condensate surface.

The initial condensate size distribution is generated based on size distributions observed experimentally. Since most PGL-3 condensates shrink and the overall PGL-3 condensate volume decreases throughout the O to E transition, the cytoplasmic PGL-3 concentration at the beginning of the simulation is expected to be below the cytoplasmic concentration that corresponds to the steady state of a condensate of an average size. Specifically, we define such steady state concentration $c_{ss}$ to be such that $\frac{d\langle R\rangle}{dt} = 0$, where $\langle R\rangle$ denotes the average radius of a given collection of condensates. From (3),

$$c_{ss} = c_{out}\left(\langle\frac{l_c(t=0)}{R(t=0)}\rangle + 1\right) . \quad (5)$$

Using this concentration as the reference, we set our initial cytoplasmic concentration to be $0.96 \times c_{ss}$.

The only remaining parameter is the kinetic rate $K$. Since variations in growth/shrinkage kinetics are clearly observed experimentally, we include these variations in the kinetic equations by having condensate specific kinetic rate. Specifically, the $i$-th condensate's kinetic rate is given by

$$K_i = \kappa\left(K + \frac{K}{2}r_i\right), \quad (6)$$

where $K$ is set to be 10 nm/s, and $r_i$ is a random number generated uniformly between $-1$ and 1. The variation in the kinetics can again be justified by the variable rugged energy landscapes in distinct condensates (fig. S10E). In addition, since MEG-3 clusters on the surface diminish a PGL-3 monomer's access to the PGL-3 condensate, the rate is reduced as a result, which is modeled by the presence of the prefactor $\kappa$, taken to be

$$\kappa(t) = (1 - 0.2)e^{-\frac{t}{\tau}} + 0.2 . \quad (7)$$

*Mutant*

For the mutant case, all parameters are identical to WT, except that since MEG-3 is absent, the capillary length returns to the expected value of $l_c = 15$ nm, and the prefactor $\kappa$ is no longer present. However, variations in the kinetic rates remain and the expressions are given by:

$$K_i = K + \frac{K}{2}r_i . \quad (8)$$

The simulation predicts a slight increase in PGL-3 volume in wild-type zygotes compared to *meg-3 meg-4* zygotes which is not observed experimentally. This suggests that MEG-3 may increase the solubility of PGL-3 slightly *in vivo*. For simplicity, we did not include this in the model.

***Modeling Pronuclear Formation (PF) to Pronuclear Migration (PM)***

*Wild-type*

Just prior to PM, condensates are quickly nucleated. MEG-3 clusters are initially found on all condensates and progressively relocalize from anterior to posterior (Fig. S8A and B). In the simulation, we assume that all posterior PGL-3 condensates are coated by MEG-3 clusters, while anterior PGL-3 condensates are not.

Due to the Pickering effect of MEG-3 clusters on the PGL-3 condensates, $l_{c,P} \ll l_{c,A}$. Specifically, we assume that $l_{c,A} = 15$ nm and $l_{c,P} = \frac{l_{c,A}}{100} = 0.15$ nm. Since the anterior and posterior condensates have distinct properties in this case, the kinetic equations are:

$$\frac{dR_{A,i}(t)}{dt} = K_{A,i}\left(\epsilon_A(t) - \frac{l_{c,A}}{R_{A,i}(t)}\right) \quad (9)$$

$$\frac{dR_{P,i}(t)}{dt} = K_{P,i}\left(\epsilon_P(t) - \frac{l_{c,P}}{R_{P,i}(t)}\right) \quad (10)$$

where the subscripts $A, P$ refer to whether the condensates are in the anterior or posterior, respectively.

Furthermore, similar the O-to-E transition, the kinetics parameters are defined as follows:

$$K_{A,i} = K + \frac{K}{2} r_{A,i} \quad (11)$$

$$K_{P,i} = \kappa\left(K + \frac{K}{2} r_{P,i}\right) \quad (12)$$

where $K = 12$ nm/sec and $r_{A,i}, r_{P,i}$ are again random numbers picked uniformly between $-1$ and 1. Compared to *meg-3 meg-4* mutants, the kinetic rate $K$ is higher in wild-type due to the fluidization effects of MEG-3 in the system (assumption 3).

In (7) and (8), the supersaturation levels $\epsilon_{A/P}$ also differ in the two sides of the cytoplasm due to the MEX-5 gradient (assumption 4). Here, we assume that $c_{A,\text{out}} = 1.2 \times c_{P,\text{out}}$ (see Eq. (2)) so that solubility of PGL-3 is higher in the anterior.

To set the initial cytoplasm PGL-3 concentration, we again first consider the steady state concentration for the average condensate size at the beginning of the simulation, which is given by

$$c_{\text{ss}} = c_{P,\text{out}} \left[ \frac{N_A}{N_{\text{tot}}} \langle \frac{l_{c,A}}{R_{A,i}(t=0)} \rangle + \frac{N_P}{N_{\text{tot}}} \langle \frac{l_{c,P}}{R_{P,i}(t=0)} \rangle + 1 \right]. \quad (13)$$

Since condensates are nucleated at the beginning of the PF stage, the cytoplasmic PGL-3 concentration is expected to be above $c_{\text{ss}}$. In our simulation, we assume that $c(t=0)$ is 10% above $c_{\text{ss}}$.

*Mutant*

In the *meg-3 meg-4* mutant, all effects of MEG-3 are absent, hence $l_{c,P} = l_{c,A} = 15$ nm, and the kinetic rate is reduced due to the absence of the accompanying fluidization effects. Specifically, the kinetic rate for the mutant is:

$$K_{\text{mutant},A,i} = K_{\text{mutant}} + \frac{K_{\text{mutant}}}{2} r_{A,i} \quad (14)$$

$$K_{\text{mutant},P,i} = K_{\text{mutant}} + \frac{K_{\text{mutant}}}{2} r_{P,i} \quad (15)$$

where $K_{\text{mutant}} = 2$ nm/sec, which is 6 times less than the WT value.

However, the MEX-5-induced spatially regulated phase separation remains and thus we take $c_{A,\text{out}}$ to be $1.2 \times c_{P,\text{out}}$.

The initial cytoplasmic far-field concentration is again assumed to 10% higher than $c_{\text{ss}}$ obtained from the initial condensate size distribution.

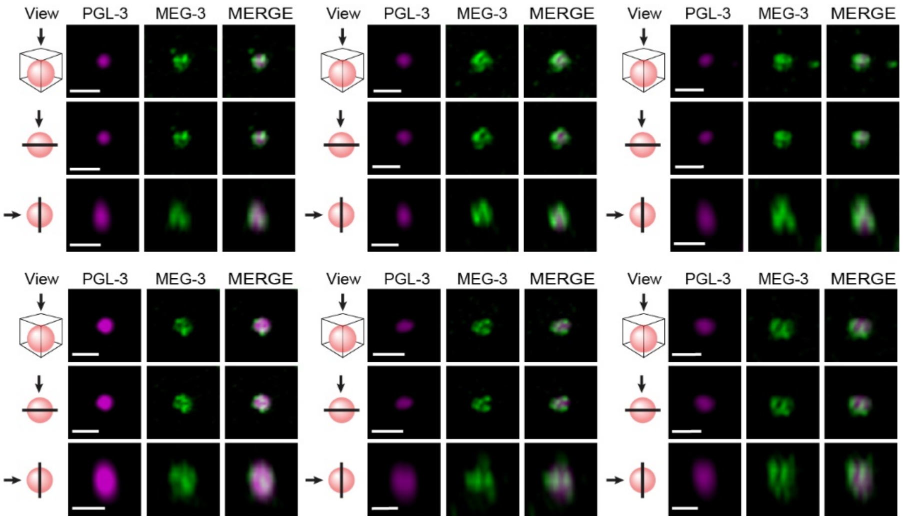


**Fig. S1. MEG-3 clusters localize to the surface of PGL-3 condensates.**

Additional examples of P granules labeled *in vivo* with MEG-3::meGFP and PGL-3::mCherry as in Fig. 1B. Top panels are a maximum projection of a Z-stack through the granule. Middle panels are a single X-Y plane through the middle of the same granule. Lower panels are a single Z-X plane through the middle of the same granule. Scale bars are 1 μm.

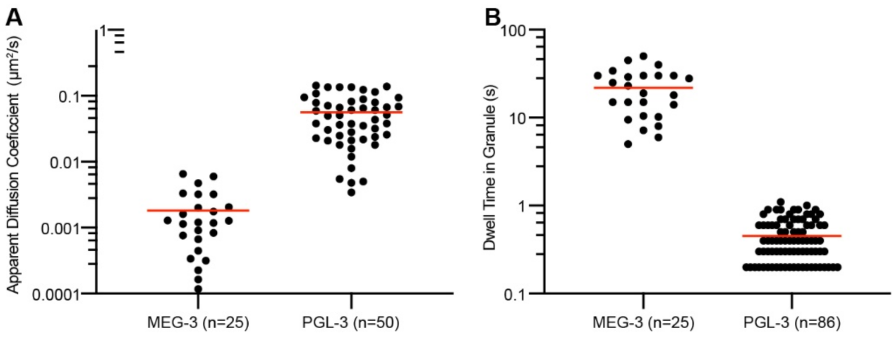


**Fig. S2. Diffusion of PGL-3 and MEG-3 in P granules.**

A) Graph depicting the apparent diffusion coefficients of PGL-3::Halo and MEG-3::Halo molecules in P granules. Each dot represents one trajectory. Red line denotes the average of the distribution.

B) Graph depicting the dwell time of PGL-3::Halo and MEG-3::Halo molecules in P granules. Each dot denotes one trajectory. Red line denotes the average of the distribution.

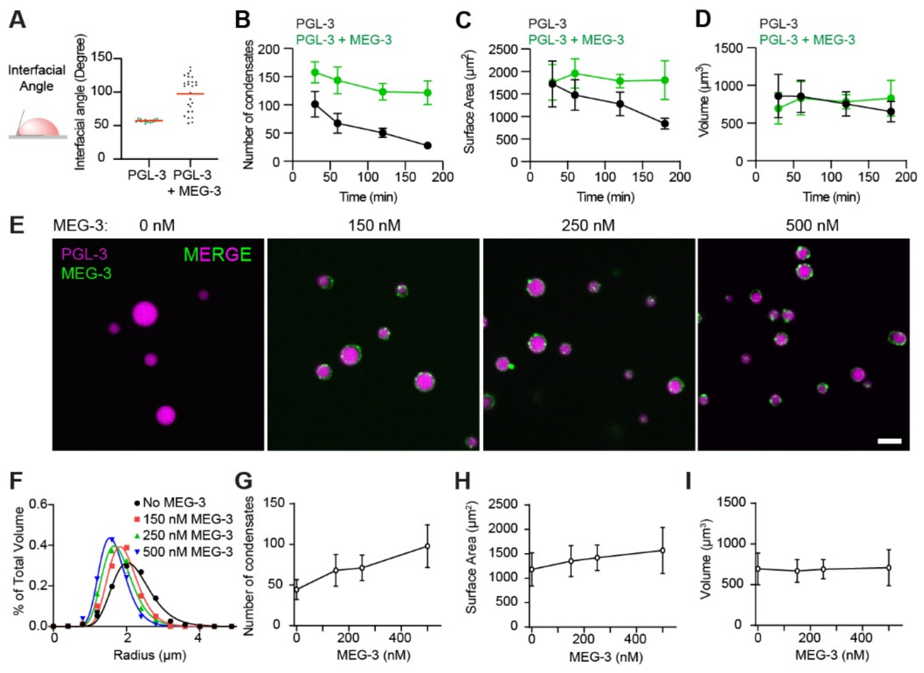


**Fig. S3. MEG-3 decreases the surface tension of PGL-3 condensates.**

A) Contact angle of PGL-3$^{488}$ condensates wetting a glass surface. Condensates were assembled as in Fig. 1E with and without MEG-3. Each dot represents a condensate. Bars represent the mean.

B-D) Graphs depicting the number (B), total surface area (C), and total volume (D) of PGL-3$^{488}$ in condensates at indicated time points with (green) and without (black) 500 nM MEG-3 assembled as in Fig. 1F. Error bars represent SD.

E) Photomicrographs of condensates of 3 μM PGL-3$^{488}$ (magenta) and 80 ng/μL *nos-2* RNA with indicated concentrations of MEG-3$^{647}$ (green) after 120 min incubation. Scale bar is 5 μm.

F) Histograms of PGL condensates assembled as in E with the indicated concentrations of MEG-3. Circles indicate the fraction of volume of PGL-3 in condensates binned by the radius of each condensate. Lines indicate fit of data to a lognormal distribution.

G-I) Graphs depicting the (G) number of condensates, (H) total surface area, (I) and total volume of PGL-3 condensates assembled as in E with the indicated concentrations of MEG-3. Error bars represent SD.

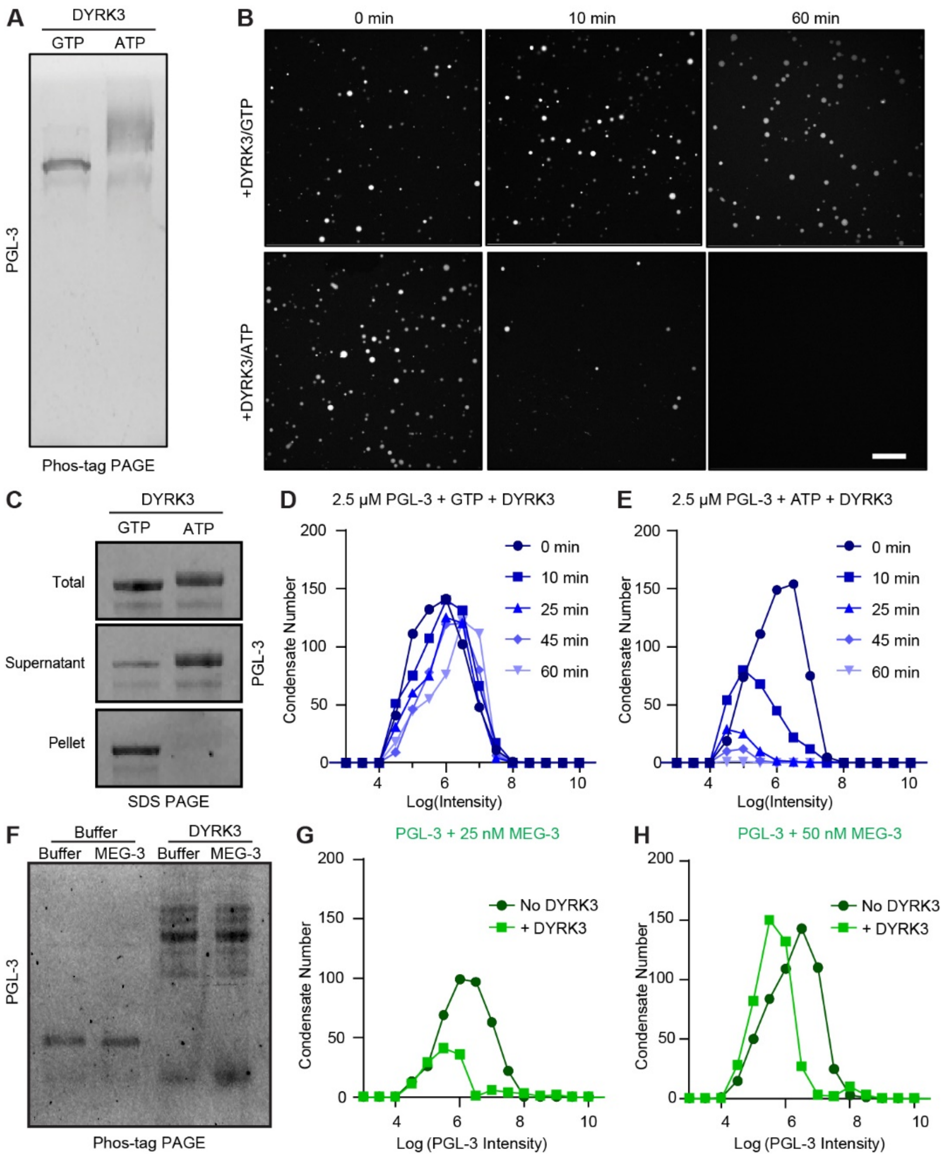


**Fig. S4. DYRK-3 phosphorylates and increases the solubility of PGL-3.**

A) Phos-tag SDS-PAGE of 2.5 µM PGL-$3^{488}$ treated with 100 nM DYRK-3 kinase with 1mM GTP or ATP. Modified PGL-3 (ATP lane) migrates more slowly than unmodified PGL-3 (GTP lane) in the Phos-tag gel.

B) Photomicrographs of a 2.5 µM PGL-$3^{488}$ emulsion at indicated time points after treatment with 100 nM DYRK-3 kinase with GTP (top panels) or ATP (bottom panels). Scale bar is 20 µM.

C) 2.5 µM PGL-$3^{488}$ treated with 100 nM DYRK3 kinase and 1mM ATP or 1mM GTP. Samples were spun at 12,000 rpm. The supernatant and pellet were collected, and samples were separated by SDS-PAGE.

D-E) Histograms of PGL condensates assembled as in (B) with GTP/DYRK3 or ATP/DYRK3 at indicated time points. Circles indicate the number of PGL-3 condensates binned by the log(Intensity) of each condensate. Colors indicate the time after addition of DYRK-3.

F) 5 µM PGL-3$^{488}$ condensates assembled with and without 100 nM MEG-3 treated with 1mM ATP and either mock kinase buffer or 100 nM DYRK3 kinase.  The reaction was stopped by addition SDS sample buffer and samples were separated by Phos-tag SDS-PAGE.
G-H) Histograms of PGL-3 condensates assembled with indicated concentrations of MEG-3 and incubated for 60 min in 100 µM ATP with and without DYRK3 – see Fig. 2D to F. Circles indicate the number of PGL-3 condensates binned by the Log(Intensity) of each condensate captured from 20 images.

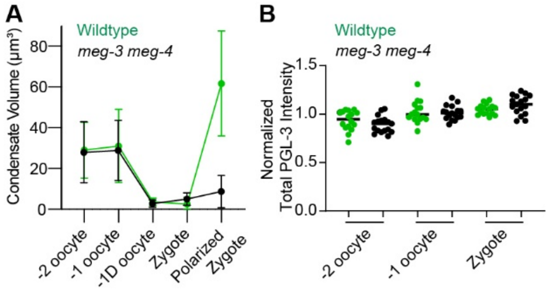


**Fig. S5. MEG-3 stabilizes the PGL-3 emulsion in fertilized zygotes.**

A) Graph depicting the total volume of PGL-3 in condensates in wild-type (green) and *meg-3 meg-4* mutants (black) during the oocyte-to-zygote transition at indicated developmental stages. Circles are the mean of ≥ 5 *in utero* images. Error bars represent SD. (The rapid rise in PGL-3 condensate volume in wild-type polarized zygotes is due to fluidization of PGL-3 condensates by MEG-3/MBK-2 – see polarization model and fig. S7).

B) Graph depicting the total PGL-3 intensity in oocytes and zygotes *in utero* represented in Fig. 3A and normalized to the average intensity of each sample. Each circle represents one oocyte/zygote in wild-type (green) or *meg-3 meg-4* (black).

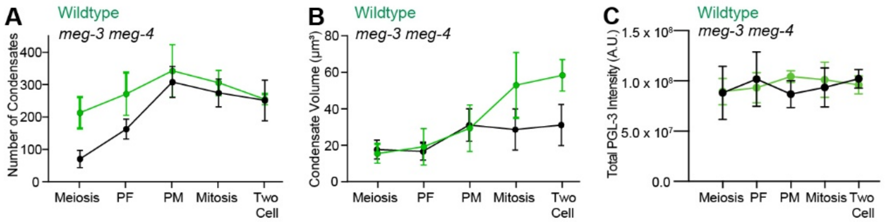


**Fig. S6. PGL-3 condensate dynamics during zygote polarization.**

A-B) Graphs showing the total number (A) or volume (B) of PGL-3 in condensates in wild-type (green) or *meg-3 meg-4* (black) zygotes at indicated developmental stages calculated from photomicrographs captured as in Fig. 4A. Circles represent the mean of ≥ 5 zygotes. Error bars represent SD.

C) Graph depicting the total PGL-3 intensity in zygotes represented in Fig. 4A. Each circle represents one zygote in wild-type (green) or *meg-3 meg-4* (black). Error bars represent SD.

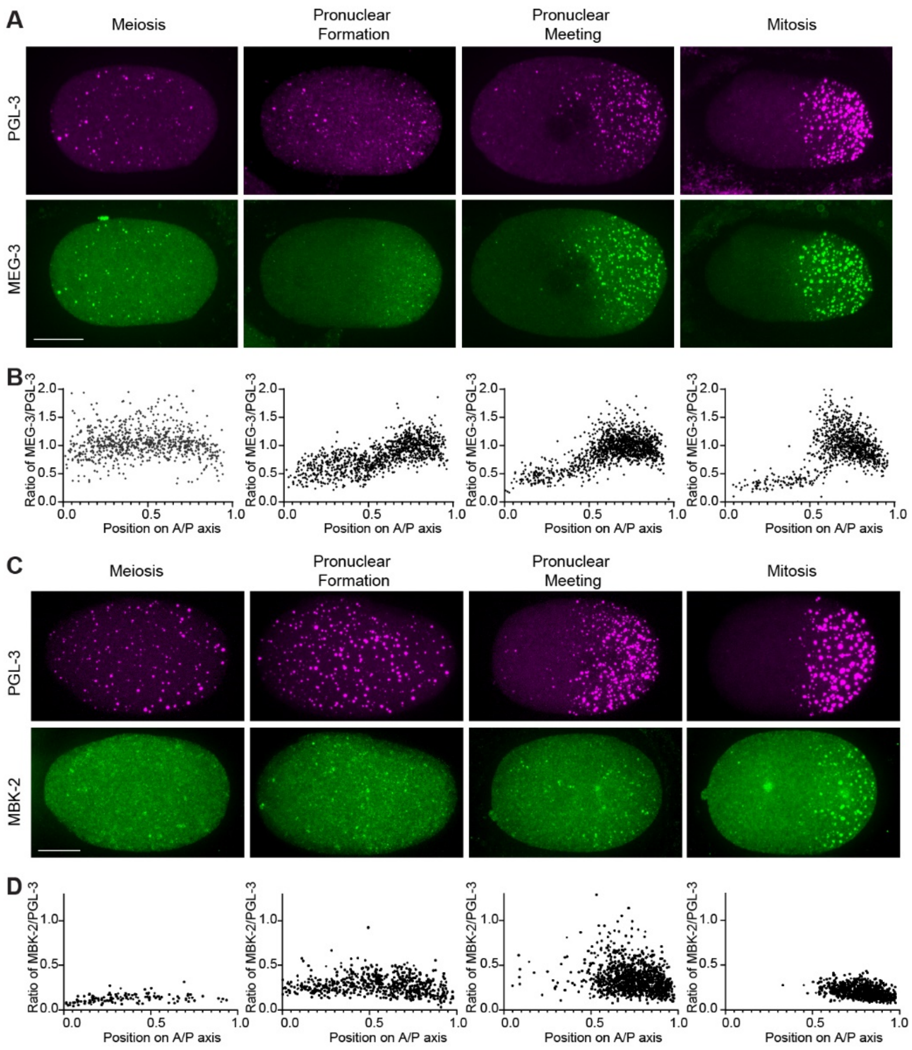


**Fig. S7. MEG-3 and MBK-2 enrichment on PGL-3 condensates.**

A) Photomicrographs of zygotes expressing MEG-3::OLLAS immunostained for OLLAS and PGL-3 at indicated developmental stages. Scale bar is 10 μm.
B) Graphs depicting the ratio of MEG-3/PGL-3 relative intensity for condensates from photomicrographs captured as in (A). Each circle represents a PGL-3 condensate plotted by position in the zygote where 0 is the anterior (A) and 1 is the posterior (P).
C) Photomicrographs of zygotes expressing PGL-3::mCherry and MBK-2::OLLAS immunostained for OLLAS at indicated developmental stages. Scale bar is 10 μm.
D) Graphs depicting the ratio of MBK-2/PGL-3 relative intensity for condensates photomicrographs captured as in (C). Each circle represents a PGL-3 condensate plotted by position in the zygote where 0 is the anterior (A) and 1 is the posterior (P).

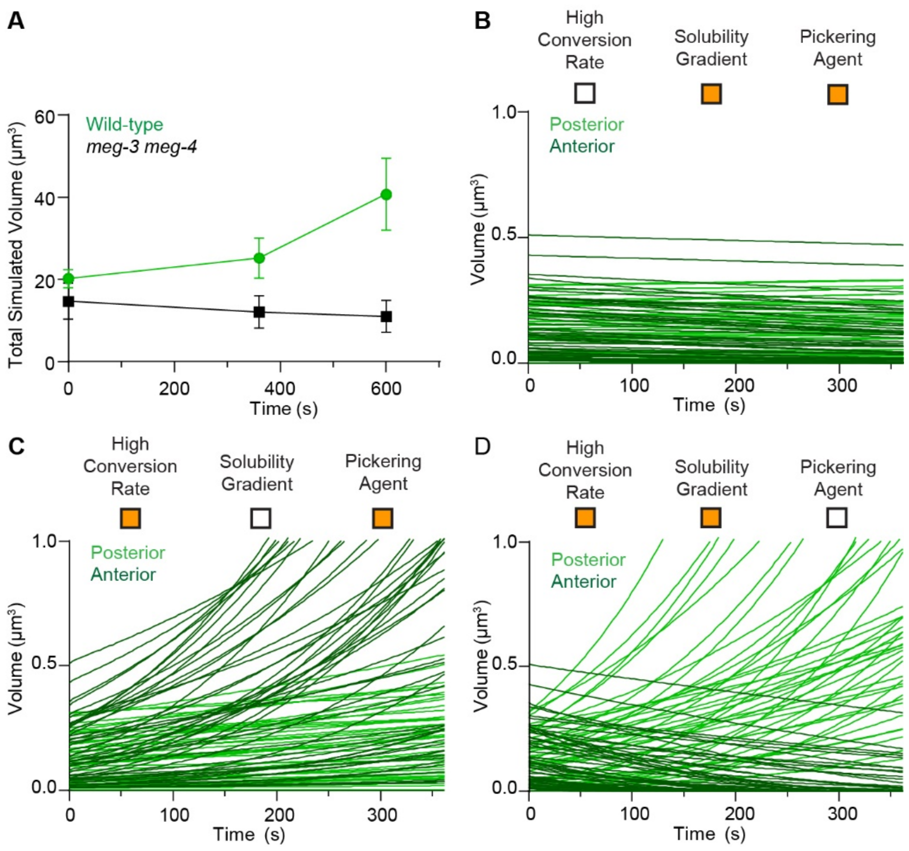


**Fig. S8.** ***In silico*** **dynamics of PGL-3 emulsion.**

A) Graph showing the simulated total volume of PGL-3 in condensates representing pronuclear formation (0s), pronuclear meeting (360s), and mitosis (600s). Compare to *in vivo* measurements shown in fig. S6B. Error bars represent SD.

B-D) Graphs showing the evolution of individual anterior (dark color) and posterior (light color) PGL-3 condensates under simulated experimental conditions as in Fig. 4L with (orange boxes) or without (white boxes) individual parameters. (B) uniformly low conversion rate (no MEG-3/MBK-2-dependent fluidization) (C) uniformly low PGL-3 solubility (no MEX-5-dependent solubility gradient) (D) uniformly high surface tension (no MEG-3 Pickering agent).

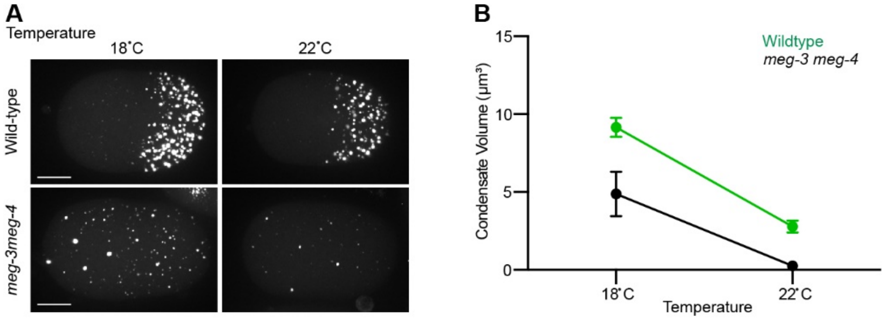


**Fig. S9. Temperature-sensitivity of the PGL-3 emulsion.**

A. Photomicrographs of PGL-3::mCherry in live zygotes (mitosis) at 18°C and after a 4 min incubation at 22°C. Scale bar is 10 μm.

B. Graph showing average PGL-3 condensate volume in zygotes (mitosis) at 18°C and after a 4 min incubation at 22°C. Error bars represent SEM.

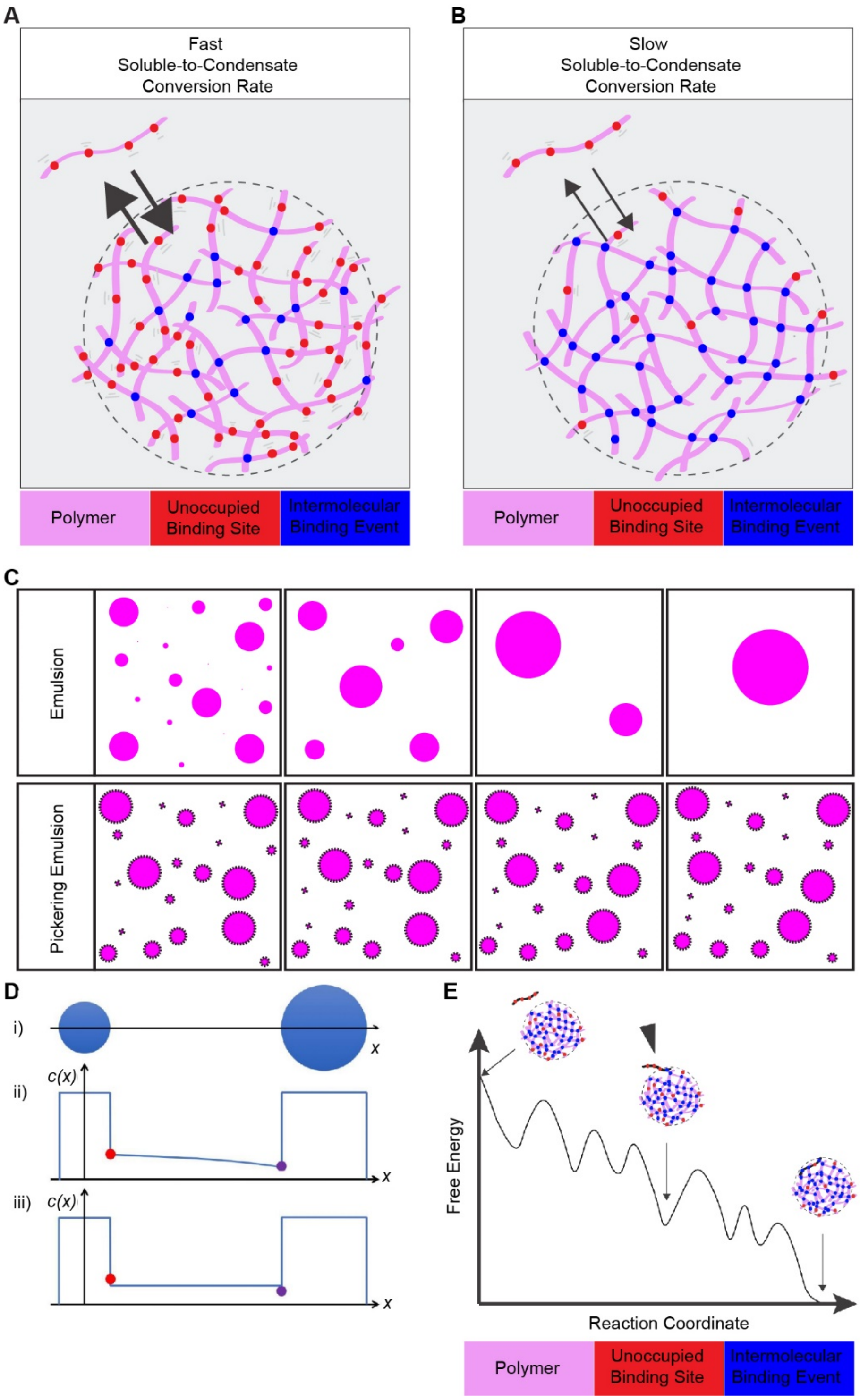

A
Fast
Soluble-to-Condensate
Conversion Rate
Polymer
Unoccupied Binding Site
Intermolecular Binding Event
B
Slow
Soluble-to-Condensate
Conversion Rate
Polymer
Unoccupied Binding Site
Intermolecular Binding Event
C
Emulsion
Pickering Emulsion
D
i)
ii)
iii)
c(x)
x
E
Free Energy
Reaction Coordinate
Polymer
Unoccupied Binding Site
Intermolecular Binding Event

**Fig. S10: Theory of condensate dynamics.**

(A-B) Schematic depictions of a protein condensate assembled from a polymer (pink) with four binding sites that can be either in a bound state (blue) or unbound state (red).
(A) In a highly dynamic (fluid) condensate, many binding sites are unbound, internal polymer rearrangement is fast and many polymers are available to bind with polymers that diffuse near the interface, leading to a high soluble-to-condensate conversion rate.
(B) In a condensate nearing kinetic arrest, many binding sites are bound, internal polymer rearrangement is slow and few polymers are available to bind polymers that diffuse near the interface, leading to a low soluble-to-condensate conversion rate.
(C) Surface tension drives emulsions to coarsen over time (top). Pickering agents (black dots) are nanoscale solids that adsorb to the surface of condensates and slow coarsening (bottom).
(D) Ostwald ripening: diffusion-limited scheme vs. conversion-limited scheme. i) Consider one small and one big condensates far away from other condensates. ii) Under the diffusion-limited scheme (the standard assumption in the Lifshitz-Slyozov theory (*60*)), the concentration profile ($x$) along the black arrow across the condensates in i) is shown in blue. In the dilute medium, the profile is obtained by solving the diffusion equation with the boundary condition set by the solute concentration right outside the two drops. Due to the Gibbs-Thomson relation (resulting from the Laplace pressure) (*61*), the smaller condensate will have a higher solute concentration (indicted by the red dot) than that of the bigger condensate (purple dot). Due to the non-zero gradient of the diffusive profile at the drop boundaries, there is an outflux of solute from the small condensate and a corresponding influx of material into the big condensate. iii) If the condensates are in the slow kinetic state (due to the rugged energy landscape leading to slow internal dynamics, fig. S10E), the rates of incorporation and exit of solutes into and out of the condensates can become limited compared to the time scale of concentration equilibration (due to diffusion) in the dilute medium. Therefore, the concentration profile in the dilute medium remains uniform, while the mismatches of the boundary condition at the condensates' boundaries lead to an outflux from the small condensate and an influx into the big condensate. These conditions represent a conversion-limited scheme.
(E) Rugged energy landscape induced slow internal dynamics. A schematic depiction of the rugged free energy of a system of one PGL-3 condensate plus one neighboring PGL-3 monomer (depicted as in A and B). The ruggedness can originate from the internal reconfiguration required, which may involve binding and unbinding of multiple protein domains, to accommodate the incorporation of the PGL-3 monomer into the condensate. A similar rugged energy landscape picture has been used to explain the experimentally observed slow rate in fibril elongation (*67*).

| *C. elegans* strains | | |
|---|---|---|
| **Strain** | **Genotype** | **Reference** |
| EGD364 | *meg-3(egx4::Halo)* | Wu et al., 2019 |
| JH3913 | *pgl-3(ax4512::Halo)* | This study |
| JH3914 | *pgl-3(ax4320::mCherry)* | This study |
| JH3918 | *hrde-1(tm1200);pgl-3(ax4320::mCherry)* | This study |
| JH3915 | *pgl-3(ax4320::mCherry);meg-3(ax3055) meg-4(ax3052)* | This study |
| JH3919 | *hrde-1(tm1200); pgl-3(ax4320::mCherry);meg-3(ax3055) meg-4(ax3052)* | This study |
| JH3916 | *mbk-2(ax4511::Ollas);pgl-3(ax4320::mCherry)* | This study |
| JH3917 | *mbk-2(ax4511::Ollas);pgl-3(ax4320::mCherry);meg-3(ax3055) meg-4(ax3052)* | This study |
| JH3920 | *pgl-3(ax4320::mCherry);meg-3(ax3054) meg-4(ax3052)* | This study |
| JH3477 | *meg-3(ax3051::Ollas) meg-4(ax3052)* | Smith et al., 2016 |
| JH3921 | *pgl-3(ax4320::mCherry);meg-3(ax4300::meGFP)* | This study |

Note: *hrde-1(tm1200)* is a mutation that suppresses the RNAi insensitivity of *meg-3 meg-4* mutants (*65*).

**Table S1. *C. elegans* strains.**

| Parameters | Values |
|---|---|
| Capillary length | 15 nm |
| Reduced capillary length due to Pickering effects, | 0.15 nm |
| Partition coefficient, | 20 |
| Cell cytoplasmic volume, | 30 pL |

Note: Capillary length estimated as described (*66*).

**Table S2. Simulation parameters.**

**Movie S1.**
*In vivo* time lapse showing PGL-3::Halo (strain JH3913). Right panel: Single molecules of PGL-3::Halo (green – $JF_{646}$ dye sparse labeling), Left panel: PGL-3::Halo labeled P granules (magenta – $JF_{549}$ dye heavy labeling) merged with single molecules of PGL-3::Halo (green – $JF_{646}$ dye sparse labeling). Images are single planes at 150ms time interval, total video time is 13.45 s, playback speed is 6.67 frames per second. Scale bar is 1 µm.

**Movie S2.**
*In vivo* time lapse showing MEG-3::Halo (strain EGD364). Right panel: Single molecules of MEG-3::Halo (green – $JF_{646}$ dye sparse labeling), Left panel: MEG-3::Halo labeled P granules (magenta – $JF_{549}$ dye heavy labeling) merged with single molecules of MEG-3::Halo (green – $JF_{646}$ dye sparse labeling). Images are single planes, total video time is 13 s, playback speed is 1 frames per second. Scale bar is 1 µm.

**Movie S3.**
*In vivo* time lapse of a wild-type 1 cell zygote from pronuclear formation to pronuclear meeting expressing PGL-3::mCherry (strain JH3914). Images are maximum intensity projections of Z planes spanning half of the depth of the zygote separated by 0.16 µm steps. Total video time is 428.4 s and playback speed is 5 frames per second. Scale bar is 5 µm.

**Movie S4.**
*In vivo* time lapse of a *meg-3 meg-4* 1 cell zygote from pronuclear formation to pronuclear meeting expressing PGL-3::mCherry (strain JH3915). Images are maximum intensity projections of Z planes spanning half of the depth of the zygote separated by 0.16 µm steps. Total video time is 456 s and playback speed is 5 frames per second. Scale bar is 5 µm.

**Movie S5.**
*In silico* simulation representing a wild-type zygote from pronuclear formation to pronuclear meeting (see Fig. 4L). The simulation includes all three model parameters: heterogeneous high conversion rate for all condensates, higher PGL-3 solubility in anterior half of the zygote, and Pickering agent on posterior condensates.

**Movie S6.**
*In silico* simulation representing a *meg-3 meg-4* zygote from pronuclear formation to pronuclear meeting (see Fig. 4M). Starting condensate sizes match range observed in *meg-3 meg-4 zygotes*. Simulation includes a uniformly low conversion rate for all condensates, higher PGL-3 solubility in anterior half of the zygote, and no Pickering agent on posterior condensates.

**Movie S7.**
*In silico* simulation representing a 1-cell zygote as in Movie S5, but with a uniformly low conversion rate (see fig. S8B)

**Movie S8.**
*In silico* simulation representing a 1-cell zygote as in Movie S5, but with uniformly low PGL-3 solubility (see fig. S8C).

**Movie S9.**
*In silico* simulation representing a 1-cell zygote as in Movie S5, but lacking the Pickering agent on posterior condensates (see fig. S8D)
